# Suprachiasmatic Neuromedin-S Neurons Regulate Arousal

**DOI:** 10.1101/2025.02.22.639648

**Authors:** Yu-Er Wu, Roberto De Luca, Rebecca Y. Broadhurst, Anne Venner, Lauren T. Sohn, Sathyajit S. Bandaru, Dana C. Schwalbe, John Campbell, Elda Arrigoni, Patrick M Fuller

**Affiliations:** Department of Neurological Surgery, University of California, Davis School of Medicine; Davis, CA 95618, USA; Department of Neurology, Division of Sleep Medicine, and Program in Neuroscience Beth Israel Deaconess Medical Center, Harvard Medical School; Boston, MA 02215, USA; Department of Biology, University of Virginia; Charlottesville, VA 22904, USA

## Abstract

Mammalian circadian rhythms, which orchestrate the daily temporal structure of biological processes, including the sleep-wake cycle, are primarily regulated by the circadian clock in the hypothalamic suprachiasmatic nucleus (SCN). The SCN clock is also implicated in providing an arousal ‘signal,’ particularly during the wake-maintenance zone (WMZ) of our biological day, essential for sustaining normal levels of wakefulness in the presence of mounting sleep pressure. Here we identify a role for SCN Neuromedin-S (SCN^NMS^) neurons in regulating the level of arousal, especially during the WMZ. We used chemogenetic and optogenetic methods to activate SCN^NMS^ neurons in vivo, which potently drove wakefulness. Fiber photometry confirmed the wake-active profile of SCN^NM^ neurons. Genetically ablating SCN^NMS^ neurons disrupted the sleep-wake cycle, reducing wakefulness during the dark period and abolished the circadian rhythm of body temperature. SCN^NMS^ neurons target the dorsomedial hypothalamic nucleus (DMH), and photostimulation of their terminals within the DMH rapidly produces arousal from sleep. Pre-synaptic inputs to SCN^NMS^ neurons were also identified, including regions known to influence SCN clock regulation. Unexpectedly, we discovered strong input from the preoptic area (POA), which itself receives substantial inhibitory input from the DMH, forming a possible arousal-promoting circuit (SCN->DMH->POA->SCN). Finally, we analyzed the transcriptional profile of SCN^NMS^ neurons via single-nuclei RNA-Seq, revealing three distinct subtypes. Our findings link molecularly-defined SCN neurons to sleep-wake patterns, body temperature rhythms, and arousal control.

**Significance Statement:** Our study’s findings provide a cellular and neurobiological understanding of how Neuromedin-S (NMS)-containing SCN neurons contribute to regulating circadian rhythms, sleep-wake patterns, body temperature, and arousal control in mammals. This research illuminates the circuit, cellular, and synaptic mechanisms through which SCN neurons regulate daily cycles of wakefulness and sleep, with implications for understanding and potentially manipulating these processes in health and disease.

## Introduction

Circadian rhythms in mammals are regulated by the hypothalamic suprachiasmatic (SCN) circadian clock. This master clock plays a central role in governing the daily temporal structure of nearly all biological processes, including the regulation of the timing and architecture of the sleep-wake cycle^1–3^. Evidence from so-called ‘forced desynchrony’ studies in both humans and animals, which involve manipulating environmental cues (such as light-dark cycles or meal timing) to assess circadian rhythms independently of external time cues, suggests that the SCN may also possess an essential role in sustaining optimal arousal levels during the ‘wake maintenance zone’ (WMZ). The WMZ occurs in the last hours of the biological day and counteracts mounting sleep pressure, thereby permitting a consolidated bout of waking.

The cellular SCN is highly heterogenous and is comprised of approximately 20,000 neurons that release over 100 identified neurotransmitters, neuropeptides, cytokines, and growth factors^4–7^. Previous research on neuropeptide-releasing neurons within the SCN has primarily focused on vasoactive intestinal polypeptide (VIP) and gastrin-releasing peptide (GRP) neurons in the ventral “core” region of the SCN, as well as arginine vasopressin (AVP) neurons in the dorsal “shell” region of the SCN^8,9^. Generally, however, selective manipulation of this different cell types has had little to no impact on level or amount of sleep or arousal^9^. Hence the specific subtypes of SCN neurons and postsynaptic targets through which the SCN regulates sleep-wake circadian rhythms, and in particular may help sustain arousal, remain elusive.

Recently, a new SCN peptide, Neuromedin S (NMS), named for its restricted expression within the SCN (SCN^NMS)^, has garnered attention for its role in circadian regulation^10,11^. Specifically, SCN^NMS^ neurons, and the Bmal1-based clock within these neurons, act as essential pacemakers in the SCN, contributing to the generation of daily rhythms in behavior. Based on this, we hypothesized that SCN^NMS^ neurons might contribute to sleep-wake regulation, in particular the regulation of arousal level, and possibly to the regulation of physiological rhythms, such as the daily body temperature (Tb) rhythm.

To test our hypothesis, we first investigated whether acute and selective activation of SCN^NMS^ neurons could trigger arousal from sleep. Additionally, we employed a genetically targeted ablation approach to selectively eliminate SCN^NMS^cells, allowing us to assess the impact of disrupting these neurons on sleep-wake activity and circadian Tb rhythms. In vivo, we used fiber photometry to evaluate state-dependent activity dynamics of SCN^NMS^ neurons. Channelrhodopsin (ChR2)-assisted circuit mapping (CRACM) provided insights into the synaptic output pathways modulating arousal state. Furthermore, modified rabies tracing helped define pre-synaptic inputs that might influence SCN^NMS^ activity. Finally, high-throughput single-nuclei sequencing (sNuc-seq) of SCN^NMS^ cells allowed us to identify distinct subtypes and characterize their molecular identities.

### Chemogenetic activation of SCN^NMS^ neurons potently drives arousal

We first aimed to determine the functional role of SCN^NMS^ neurons in arousal control through conditional chemogenetic activation using the excitatory Designer Receptor Exclusively Activated by a Designer Drug (DREADD; AAV-hSyn-DIO-hM3Dq, hereafter referred to as AAV-hM3Dq). We performed bilateral injections of AAV-hM3Dq into the SCN of n=7 NMS-IRES-Cre (NMSiCre) mice (Fig. 1a). Three weeks later we administered the hM3Dq ligand, clozapine-N-oxide (CNO, 0.3 mg/kg, intraperitoneal) to these mice. CNO injections consistently produced robust c-Fos expression in hM3Dq(+) SCN^NMS^ neurons (Fig. 1b and Extended Data Fig. 1). Subsequently, we monitored the sleep-wake patterns and arousal states of the mice based on both EEG and EMG signals. Following injections of CNO at 10 AM (a time approximating peak sleep drive in mice), wakefulness significantly increased during the first hour. Specifically, wakefulness reached 82% ± 2.5% or 87.8% ± 4.5% after 0.3 mg/kg or 1 mg/kg CNO, respectively, which was significantly higher than wakefulness after saline administration. Concurrently, NREM sleep decreased significantly during the first hour after CNO, with only 16.9% ± 2.6% (0.3 mg/kg CNO) or 12.2% ± 4.5% (1 mg/kg CNO) of time spent in NREM sleep. Additionally, REM sleep was completely suppressed during the first 3 hours after CNO injection (Fig. 1c,d).

**Fig. 1.**
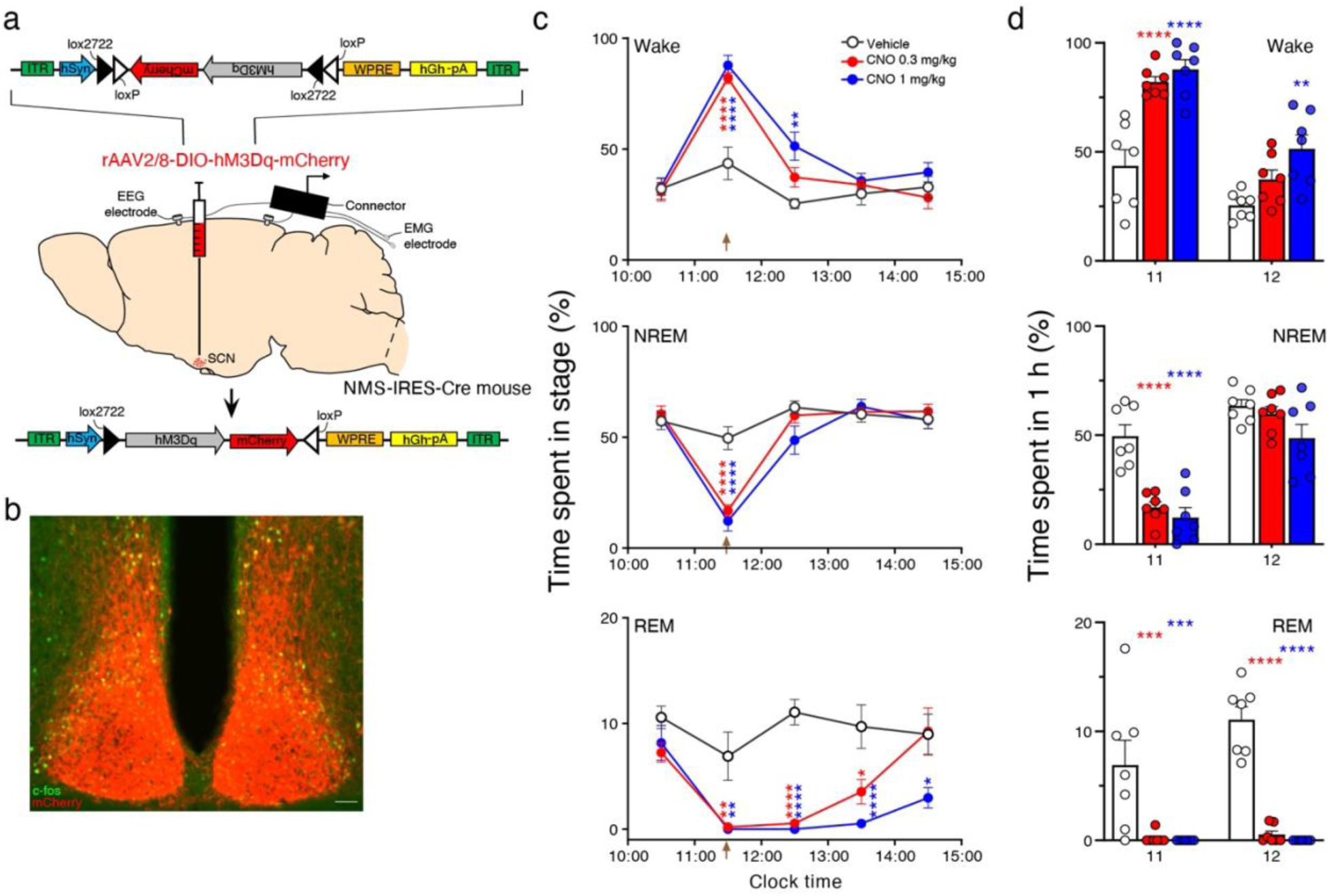
Chemogenetic activation of SCN^NMS^ neurons increases wakefulness. **a**, Schematic illustration of the experimental setup: AAV-hSyn-DIO-hM3Dq-mCherry was bilaterally injected into the SCN of NMS-IRES-Cre mice. Two weeks later, the mice were equipped with an EEG/EMG headset to record sleep-wake patterns. **b**, Photomicrograph depicting immunofluorescent expression of hM3Dq-mCherry (Red) and cFos (green) in SCN^NMS^ neurons. Scale bar, 200μm. Clear co-localization of mCherry expression and cFos induction, indicating activation of hM3Dq-expressing neurons in response to CNO. **c**, Time course of changes in wakefulness, NREM sleep, and REM sleep following the administration of vehicle or CNO in mice expressing hM3Dq in SCN^NMS^ neurons. Each circle represents the mean hourly amount of time spent in each stage. Values are the mean ± SEM for each group (n = 7 mice). Two-way ANOVA followed by Sidak post hoc test is shown. *p < 0.05; **p < 0.01; ****p < 0.0001. **d,** Total time spent in wakefulness, NREM sleep, and REM sleep during the 1^st^ h and 2^nd^ h recording following the vehicle or CNO administration (n = 7 mice). Two-way ANOVA followed by Sidak post hoc test is shown. **p < 0.01; ***p < 0.001; ****p < 0.0001.

### Selective ablation of SCN^NMS^ neurons and sleep-wake

We next sought to investigate whether SCN^NMS^ neurons play a crucial role in establishing and maintaining the sleep-wake cycle. To do so, we administered bilateral injections of an AAV vector expressing a Cre-dependent diphtheria toxin (AAV-FLEX-DTA-AAV) into the SCN of n=5 NMSiCre mice and n=7 WT mice (referred to as SCN^NMS-DTA^ and SCN^WT-DTA^; see Fig. 2a,b). The AAV-FLEX-mCherry-DTA vector expresses diphtheria toxin A in neurons containing Cre recombinase and mCherry in neurons lacking Cre expression (the latter allows us to identify the injection site). We conducted electroencephalogram (EEG) recording and analysis on SCN^NMS-DTA^ and SCN^WT-DTA^ mice to assess the impact of SCN^NMS^ cell loss on sleep-wake circadian rhythms (see Fig. 2c). Compared to control SCN^WT-DTA^ mice, SCN^NMS-DTA^ mice exhibited a significant decrease in wakefulness during the dark (active) period in the light-dark cycle (LD; as shown in Figs. 2c,d). Notably, this reduction occurred during the last 3 hours of the active period, a time of mounting sleep drive/pressure in mice and humans (98.6 ± 3.3 minutes vs. 124.4 ± 6.8 minutes wakefulness in control mice). The reduced wakefulness in SCN^NMS-DTA^ mice was replaced by increased amounts of both NREM sleep and REM sleep during this period (see Fig. 2e). In summary, our findings highlight the essential role of SCN^NMS^ neurons in maintaining normal arousal levels, particularly during the later hours of the active period, i.e., the wake maintenance zone, WMZ.

**Fig. 2.**
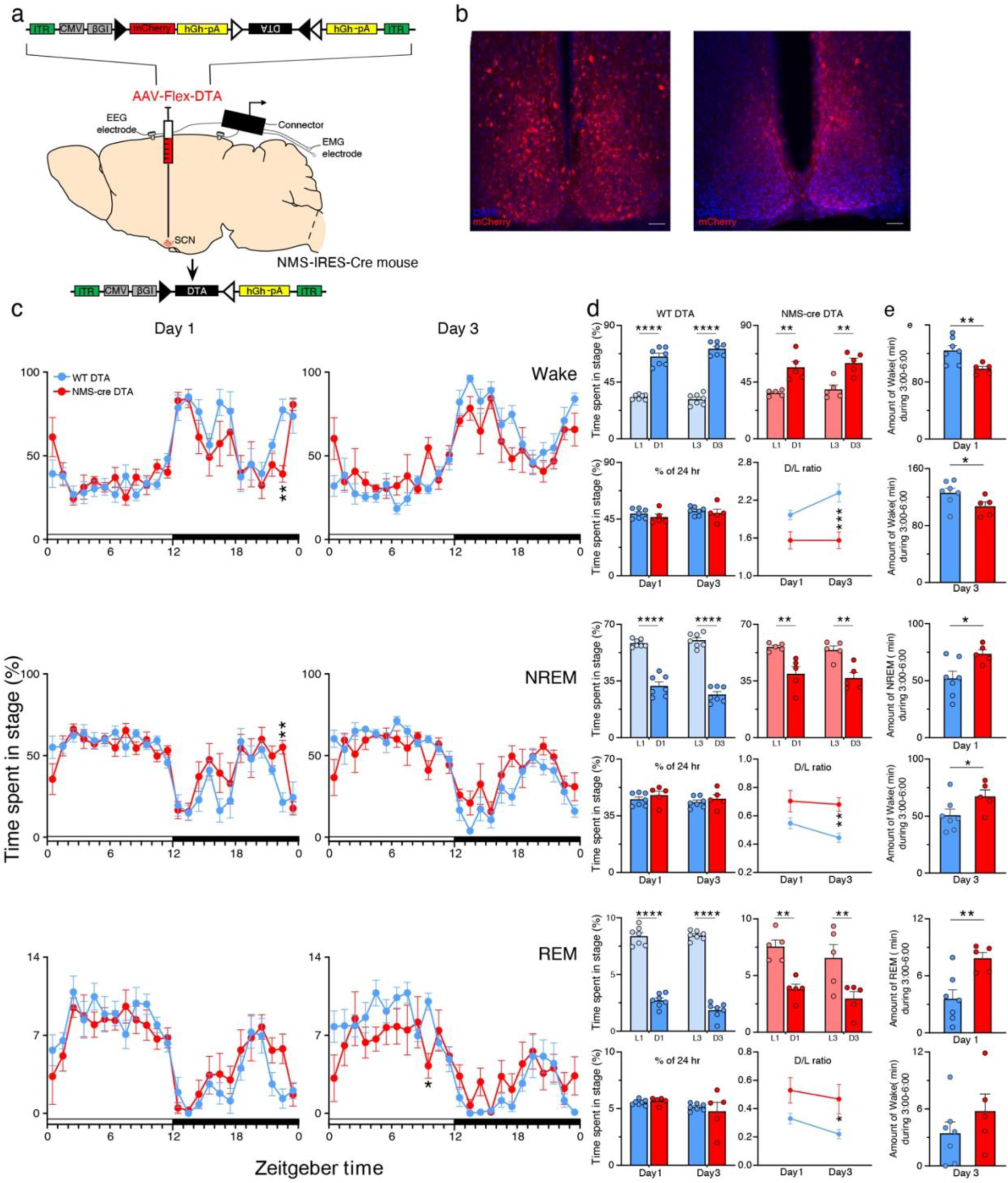
Effect of selective ablation of SCN^NMS^ neurons on sleep–wake. **a**, Experimental schematic: AAV-CMV-Flex-DTA-mCherry was bilaterally injected into the SCN of NMS-IRES-Cre mice. Mice were subsequently implanted with an EEG/EMG electrode to record sleep-wake. **b**, Photomicrograph depicting expression of Flex-DTA-mCherry (Red) in SCN^NMS^ neurons in NMS-IRES-Cre mice or wild-type littermates (WT) mice. As non-NMS neurons do not express Cre and therefore are not killed by the Cre-dependent AAV-DTA, these neurons expressing mCherry indicate the injection site. Scale bar, 200μm. **c**, Hourly distribution of wakefulness, NREM sleep, and REM sleep for two days. Bars indicate mean ± SEM for the group (n = 5 to 7 mice). Two-way ANOVA followed by Sidak post hoc test is shown. *p < 0.05; **p < 0.01. **d**, Upper panels: Amounts (mean ± SEM) of the vigilance stages during the two days light (L1, L3) and dark (D1, D3) periods in WT mice (upper left panels; n = 7 mice) and NMS-IRES-cre mice (upper right panels; n = 5 mice). Lower panels: 24 h amounts (mean ± SEM) of the vigilance stages during the two days (Day1, Day3; lower left panels) and dark to light (D/L) ratio for each vigilance stage stages during the two days Day1, Day3; lower right panels). Two-way ANOVA followed by Sidak post hoc test is shown. *p < 0.05; **p < 0.01; ***p < 0.001; ****p < 0.0001. **e**, Total time spent in wakefulness, NREM sleep, and REM sleep during the 3:00 am to 6:00 am (n = 5 to 7 mice). Paired two-tailed t-test. *p < 0.05; **p < 0.01.

### Selective ablation of SCN^NMS^ neurons produces arrythmicity of the body temperature (Tb) rhythm

To investigate the impact of SCN^NMS^ cell loss on circadian rhythms of body temperature (Tb), we administered AAV-DTA injections into the SCN of NMSiCre mice. These mice (SCN^NMS-DTA^) were also instrumented for chronic Tb recordings (see Fig. 3). After three weeks in a 12:12 light-dark cycle, the mice were released into constant darkness (DD) conditions for 10 days. As expected, in DD, control mice (SCN^WT-DTA^) displayed high-amplitude free-running rhythms in Tb, whereas SCN^NMS-DTA^ mice became almost immediately arrhythmic (as shown in Fig. 3a,b). Therefore, our findings demonstrate that SCN^NMS^ neurons play a crucial role in maintaining coherent circadian rhythms in Tb.

**Fig. 3.**
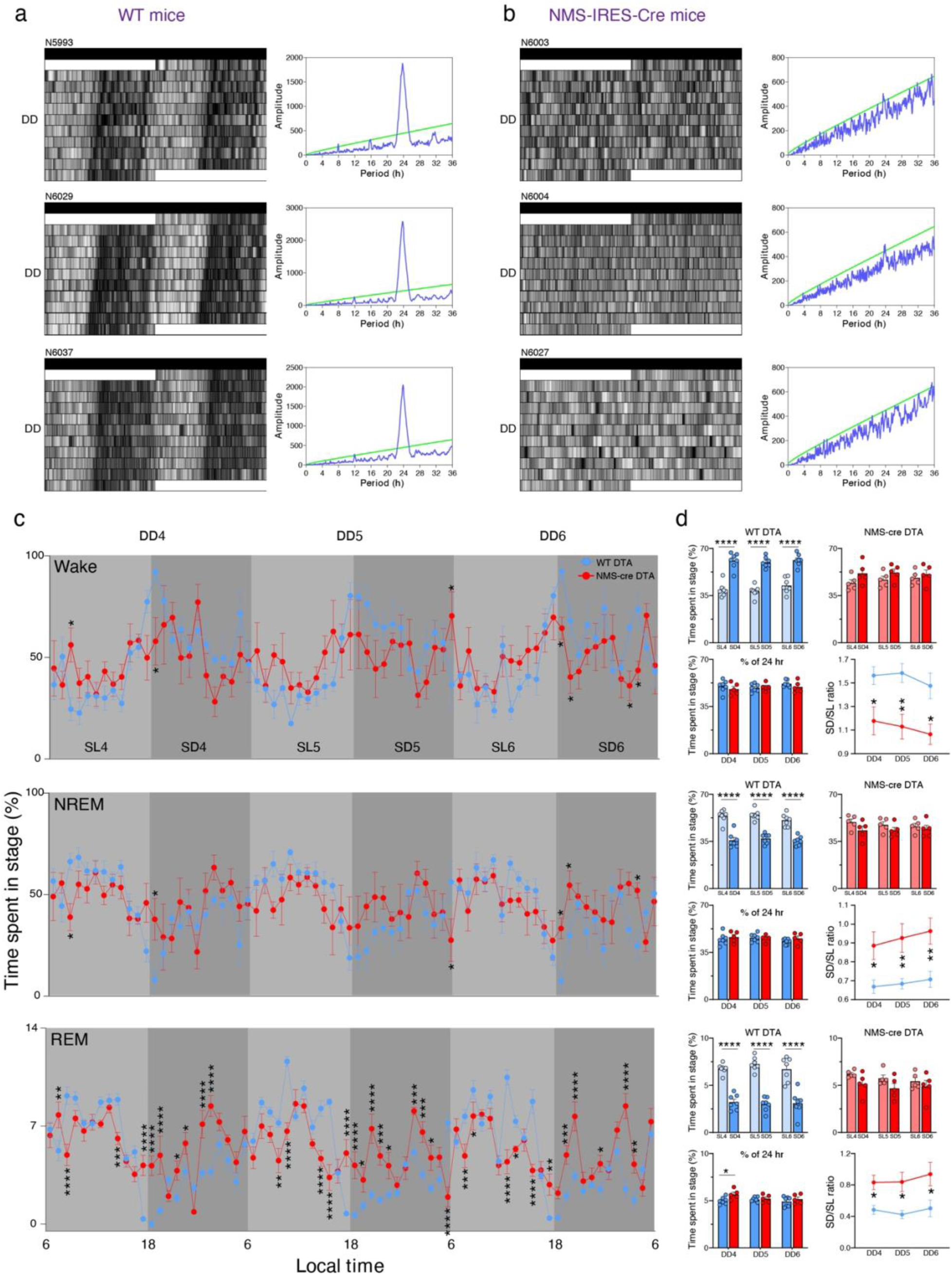
Circadian Rhythm of Body Temperature is Arrhythmic following Selective Ablation of SCN^NMS^ Neurons. **a,b**, Representative actograms showing body temperature (Tb) circadian rhythms (gray scale: darker represents higher temperature) and associated periodograms during ten days in constant dark (DD) from three WT mice (a) and three NMS-IRES-Cre mice (b). Note a profound disruption of Tb rhythms in NMS-IRES-Cre mice, while these rhythms remain unaltered in WT mice. **c**, Time-course of wakefulness, NREM sleep, and REM sleep during the 4^th^, 5^th^ and 6^th^ day in constant darkness. Bars indicate mean ± SEM for the group (n = 5 to 7 mice). Two-way ANOVA followed by Sidak post hoc test is shown. *p < 0.05; **p < 0.01; ***p < 0.001; ****p < 0.0001. **d**, Upper panels: Amounts (mean ± SEM) of the vigilance stages during the 4^th^, 5^th^ and 6^th^ subjective light (SL4, SL5, SL6) and subjective dark (SD4, SD5, SD6) periods in WT mice (upper left panels; n = 7 mice) and NMS-IRES-cre mice (upper right panels; n = 5 mice). Lower panels: 24 h amounts (mean ± SEM) of the vigilance stages during the 4^th^, 5^th^ and 6^th^ day in constant darkness (DD4, DD5, DD6; lower left panels) and subjective dark to subjective light (D/L) ratio for each vigilance stage stages during the 4^th^, 5^th^ and 6^th^ day in constant darkness (DD4, DD5, DD6; lower right panels). Two-way ANOVA followed by Sidak post hoc test is shown. *p < 0.05; **p < 0.01; ****p < 0.0001.

### Optogenetic stimulation of SCN^NMS^ neurons rapidly triggers arousal

As chemogenetic activation of SCN^NMS^ neurons was effective in inducing changes in wakefulness, we next sought to determine whether optogenetic activation of this population would be sufficient to promote wakefulness transitions from both NREM sleep and REM sleep in vivo. To do so, we stereotaxically injected AAVs expressing channelrhodopsin-2 (AAV10-EF1a-DIO-hChR2(H134R)-eYFP, hereafter referred to as AAV-ChR2) bilaterally into the SCN of NMSiCre mice and their wild-type (WT) littermates (control mice). We then installed a headset capable of optogenetically stimulating SCN^NMS^ neurons while concurrently recording the electroencephalogram/electromyogram (EEG/EMG) (Fig. 4a-c).

**Fig. 4.**
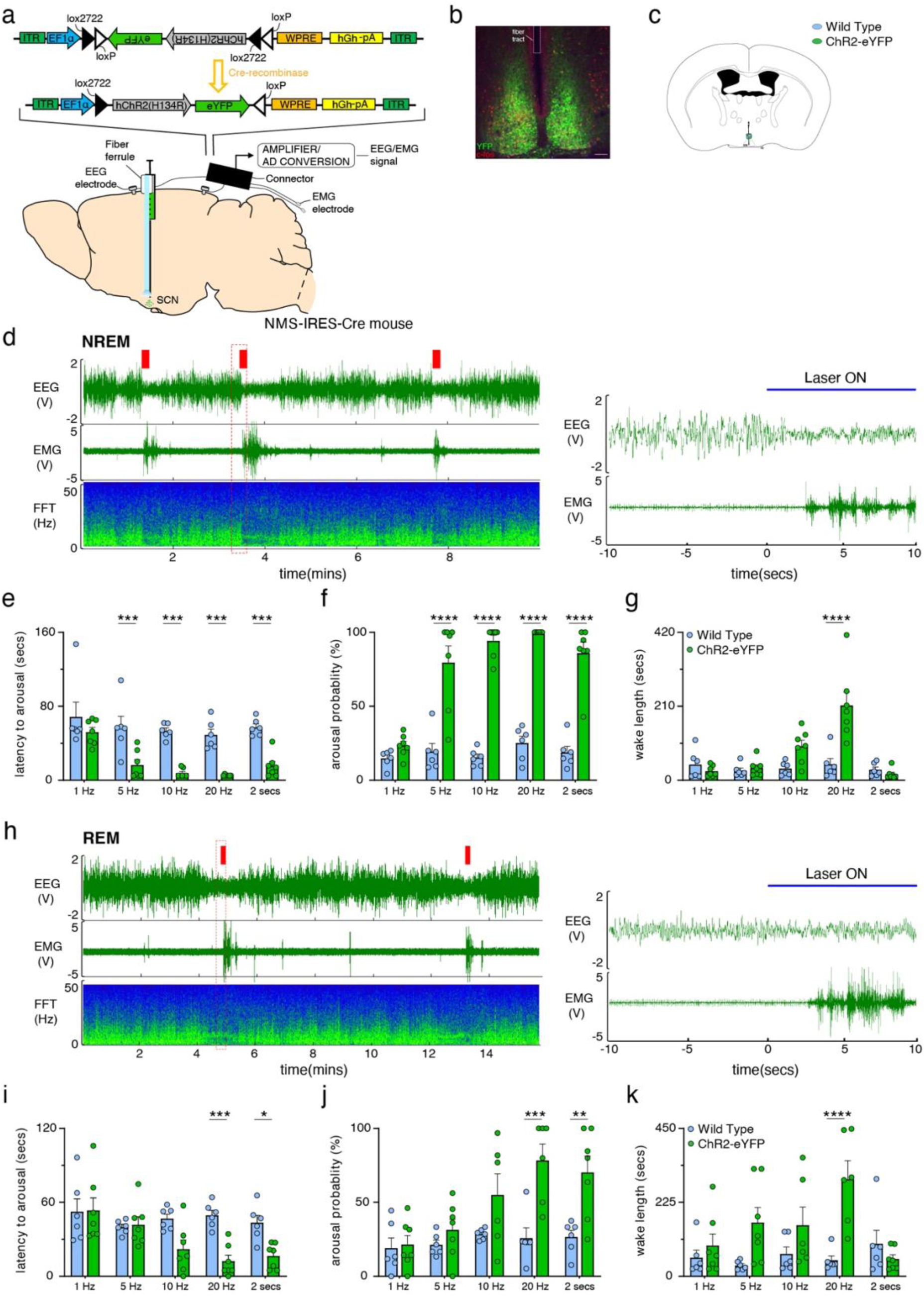
Optogenetic Stimulation of SCN^NMS^ Neurons in vivo Rapidly Triggers Arousal. **a**, Experimental schematic: AAV-EF1a-DIO-ChR2-eYFP was bilaterally injected into the SCN of NMS-IRES-Cre mice and an optical fiber was implanted over the injection site. Mice were also equipped with an EEG/EMG headset to record sleep-wake. **b**, Photomicrograph depicting expression of ChR2-eYFP in SCN^NMS^ neurons (green) and optic fiber placement. Scale bar, 200μm. **c**, Location of the optic fiber tips within the SCN of ChR2-eYFP-expressing (green) and control (blue) mice. **d**, Left: example EEG/EMG traces and corresponding FFT analysis from a mouse expressing ChR2-eYFP in the SCN following optogenetic stimulation of SCN^NMS^ cell bodies during NREM sleep (red bars; 10 s, 20 Hz, 10-ms pulses). Right: higher magnification of the red boxed area in the left-hand panel. Following optogenetic stimulation onset, the mouse awoke almost immediately. **e,f,g**, Latency to arousal (e), arousal probability during the optogenetic stimulation (f), and length of the wake episode following the stimulation (g) in wild-type (WT) and ChR2-eYFP mice over a range of stimulation frequencies delivered during NREM sleep. Bars represent mean ± SEM for the group (n = 6 to 7 mice per stimulation frequency per genotype), and individual data points represent the mean values for individual mice (calculated from 20 stimulations per stimulation frequency). **h**, Left: example sleep-wake recording from a mouse expressing ChR2-eYFP in the SCN following optogenetic stimulation of SCN^NMS^ cell bodies during REM sleep (red bars; 10 s, 20 Hz, 10-ms pulses). Right: higher magnification of the red boxed area in the left-hand panel is shown. **i,j,k**, Latency to arousal (i), arousal probability during the optogenetic stimulation (j), and length of the wake episode following the stimulus (k) in WT and ChR2-eYFP mice over a range of stimulation frequencies delivered during REM sleep. Bars represent mean ± SEM for the group (n = 6 to 7 mice per stimulation frequency per genotype), and individual data points represent the mean values for individual mice (calculated from 20 stimulations per stimulation frequency). Two-way ANOVA followed by Sidak post hoc test is shown. *p < 0.05; **p < 0.01; ***p < 0.001; ****p < 0.0001.

As expected, the injections of AAV-ChR2 resulted in robust and specific expression of ChR2-eYFP in SCN^NMS^ neurons (Fig. 4b), while the AAV-ChR2 injections in wildtype (WT) littermates did not yield any expression of eYFP. Using online sleep detection (OSD) as previously described^12^, we determined when the mice entered NREM sleep or REM sleep. During the light phase, we delivered brief (10 s, 1 Hz, 5Hz, 10 Hz, 20 Hz) 473-nm blue light stimulations to SCN^NMS^ cell bodies through the implanted optical fiber, specifically during either NREM or REM sleep. Notably, 5Hz, 10 Hz and 20 Hz of optical activation of SCN^NMS^ cell bodies reliably induced transitions to wakefulness with a short latency of less than 2 s from NREM sleep (Fig. 4d,e). The probability of inducing a short latency arousal increased with increasing light pulse stimulation frequencies (Fig. 4f).

Although the short 10-second optogenetic stimulation was sufficient to rouse the mice from NREM sleep, it was observed the effect of the stimulation at 5 Hz and 10 Hz was typically short-lasting, and the resulting wake episodes following stimulation showed comparable durations between ChR2-eYFP-expressing mice and control mice (Fig. 4g). However, when the stimulation frequency was increased to 20 Hz, a notable and significant increase in the duration of wake bouts was observed. In contrast to stimulation delivered during NREM sleep, high-frequency stimulations of SCN^NMS^ neurons were also sufficient to induce arousals from REM sleep, whereas low-frequency stimulations did not yield the same response (Fig. 4h-k). These findings demonstrate that acute activation of SCN^NMS^ neurons through optogenetics drives wakefulness in mice.

### Activation of SCN^NMS^ terminals within the dorsomedial hypothalamus (DMH) promote arousal

The foregoing work also revealed that SCN^NMS^ neurons provide dense innervation of the DMH, i.e., ChR2 anterogradely traffics to terminals. We therefore investigated whether activation of SCN^NMS^ terminals in the DMH would be sufficient to rouse mice from NREM sleep. To achieve this, we expressed ChR2-eYFP in SCN^NMS^ neurons and selectively stimulated SCN^NMS^ terminals in the DMH using optic fibers (Fig. 5a-c). Our results demonstrated that photoactivation of SCN^NMS^ terminals reliably induced transitions to wakefulness after a short latency (<2 s) from NREM sleep (Fig. 5d,e). Moreover, the probability of inducing a short latency arousal increased with higher frequencies of light pulse stimulation (Fig. 5f). In contrast to stimulation delivered during NREM sleep, 20 Hz frequency stimulations of SCN^NMS^ neurons were also sufficient to induce arousals from REM sleep, whereas other stimulations did not yield the same response (Fig. 5h-k). Consistent with the activation of SCN^NMS^ cell bodies, optogenetic stimulation of SCN^NMS^ terminals in the DMH effectively induced arousal from NREM sleep.

**Fig. 5.**
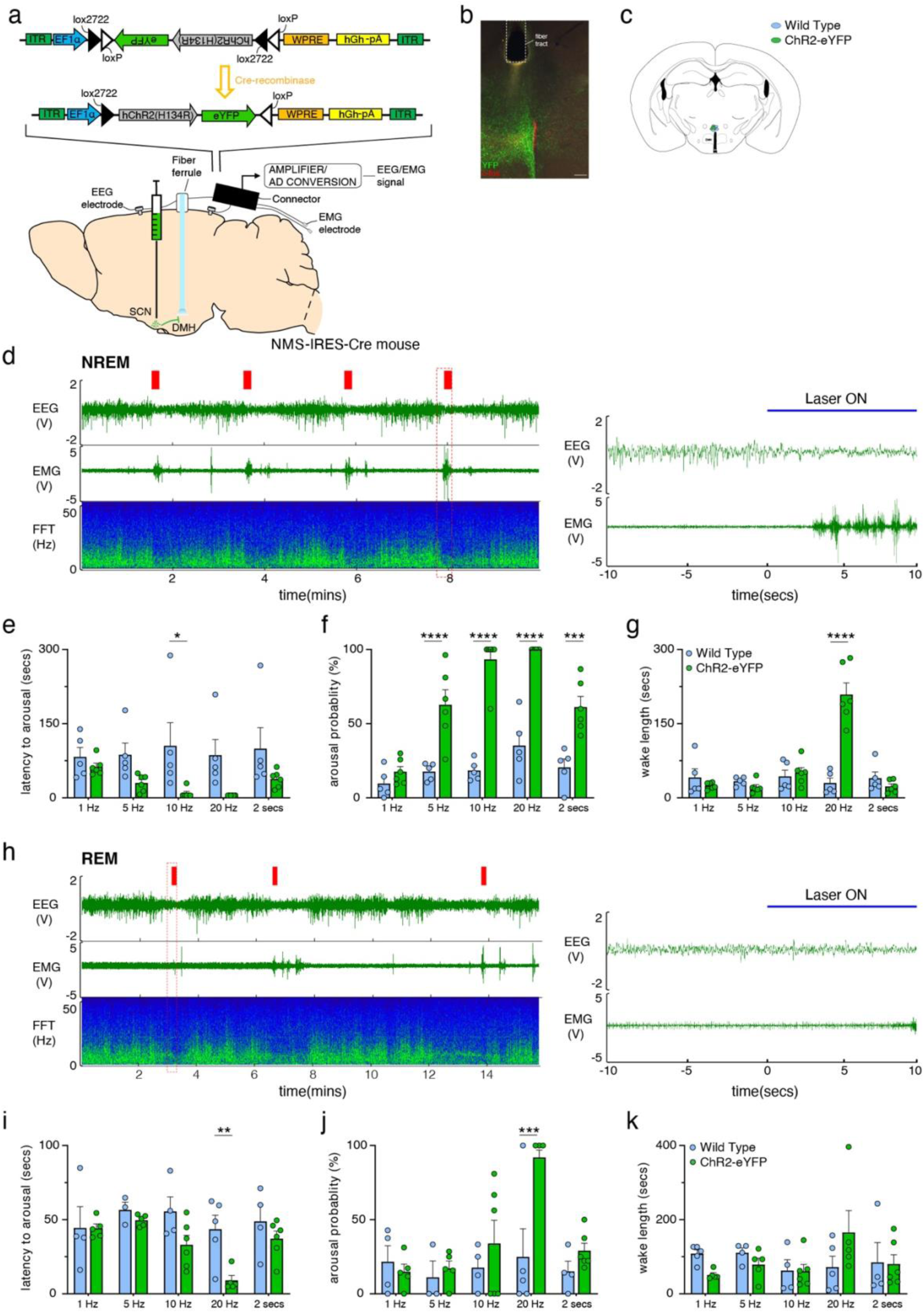
Optogenetic stimulation of SCN^NMS^ terminals in the DMH area rapidly triggers arousal. **a**, Experimental schematic: AAV-EF1a-DIO-ChR2-eYFP was bilaterally injected into the SCN of NMS-IRES-Cre mice and an optical fiber was implanted over the ipsilateral DMH area. Mice were also equipped with an EEG/EMG headset to record sleep-wake. **b**, Photomicrograph depicting expression of ChR2-eYFP expression in SCN^NMS^ terminals in the DMH area (green) and optic fiber placement. Scale bar, 200μm. **c**, Location of the optic fiber tips within the DMH area of ChR2-eYFP-expressing (green) and control (blue) mice. **d**, Left: Typical example polygraphic recording from a mouse expressing ChR2-eYFP in the SCN following optogenetic stimulation of SCN^NMS^/DMH axon terminals during NREM sleep (red bars; 10 s, 20 Hz, 10-ms pulses). Right: higher magnification of the red boxed area in the left-hand panel is shown. Following optogenetic stimulation onset, the mouse awoke almost immediately. **e,f,g**, Latency to arousal (e), arousal probability during the optogenetic stimulation (f), and length of the wake episode following the stimulation (g) in WT and ChR2-eYFP mice over a range of stimulation frequencies delivered during NREM sleep. Bars represent mean ± SEM for the group (n = 5–6 mice per stimulation frequency per genotype), and individual data points represent the mean values for individual mice (calculated from 20 stimulations per stimulation frequency). **h**, Left: example sleep-wake recording from a mouse expressing ChR2-eYFP in the DMH following optogenetic stimulation of SCN^NMS^/DMH axon terminals during REM sleep (red bars; 10 s, 20 Hz, 10-ms pulses). Right: higher magnification of the red boxed area in the left-hand panel is shown. **i,j,k**, Latency to arousal (i), arousal probability during the optogenetic stimulation (j), and length of the wake episode following the stimulus (k) in WT and ChR2-eYFP mice over a range of stimulation frequencies delivered during REM sleep. Bars represent mean ± SEM for the group (n = 5 to 6 mice per stimulation frequency per genotype), and individual data points represent the mean values for individual mice (calculated from 20 stimulations per stimulation frequency). Two-way ANOVA followed by Sidak post hoc test is shown. *p < 0.05; **p < 0.01; ***p < 0.001; ****p < 0.0001.

### Confirming functional connectivity between SCN^NMS^ and DMH neurons via CRACM

To validate synaptic connectivity between SCN^NMS^ neurons and neurons within the dorsomedial hypothalamus (DMH), channelrhodopsin-2 (ChR2)-assisted circuit mapping (CRACM) was performed on brain slices. An adeno-associated virus (AAV) conditionally expressing ChR2 (AAV10-EF1a-FLEX-hChR2(H134R)-eYFP) was unilaterally injected into the SCN of NMSiCre mice. Brain slices containing the DMH were prepared, and whole-cell patch-clamp recordings were performed on individual DMH neurons while photostimulating the SCN^NMS^ terminals within the DMH (Fig. 6a,b). Photostimulation evoked inhibitory postsynaptic currents (IPSCs) in DMH neurons (11 out of 22), confirming functional synaptic connectivity. The average peak amplitude of photo-evoked IPSCs was 71.41±31.41 pA (n=15). The latency of photo-evoked IPSCs was 5.71±0.42 ms (n=15) (Fig. 6c-f). These results confirm that SCN^NMS^ neurons form functional inhibitory synapses onto DMH neurons, validating monosynaptic connectivity between these neuronal populations. We also expressed ChR2 in VGat expressing neurons of the DMH which resulted in dense ChR2 labelled fibers in VLPO. Photostimution of the DMH GABA input in VLPO in brain slices produced opto-evoked GABAergic IPSCs in VLPO neurons (Fig. 6g) suggesting that SCN^NMS^ neuron arousal responses could be mediated via the DMH^GABA^-VLPO circuit. To test this possibility, we expressed ChR2 in SCN^NMS^ neurons and placed a second injection of fluorescent cholera toxin B (F-CTB) into the VLPO (Fig. 6h,i). We then recorded from VLPO DMH projecting neurons while photostimulating the SCN^NMS^ input. Interestingly, of the SCN^NMS^ input was directed to DMH neurons that did not project to the VLPO (Fig. 6k,j), suggesting that SCN^NMS^ neurons could promote arousal by disinhibiting a DMH^GABA^ input to the VLPO through a GABAergic circuit within the DMH.

**Fig. 6.**
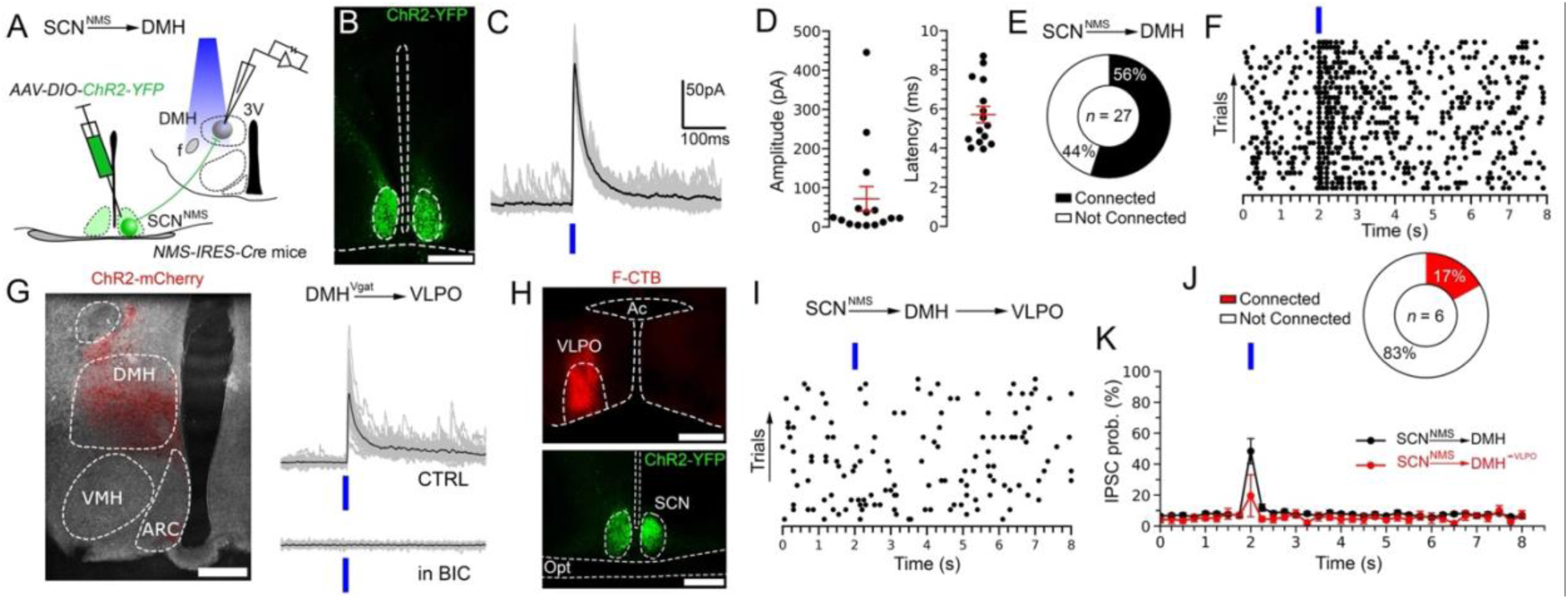
SCN^NMS^ → DMH → VLPO circuit. **A,** Schematic of the experimental design used to map connectivity between SCN^NMS^ neurons and DMH neurons (SCN^NMS^→DMH). AAV10-EF1a-DIO-hChR2(H134R)-eYFP was injected bilaterally into the SCN of NMS-IRES-Cre mice, and recordings were made from the DMH in brain slices while photostimulating the SCN^NMS^ axon terminals. **B**, Image of a SCN^NMS^ neurons expressing ChR2-eYFP (green). (Scale bar, 500 µm). **C**, Photo-evoked IPSCs from a DMH neuron recorded in whole-cell mode during photostimulation of SCN^NMS^→DMH axon terminals. 30 individual IPSCs (grey) and average IPSC (black). **D**, Mean photo-evoked IPSC amplitude (left) and latency (right) recorded from DMH neurons (n = 15; mean ±SEM in red). **E**, Percentages of the DMH neurons in which photostimulation of the SCN^NMS^ input produced opto-evoked IPSC (connected). Total number of DMH recorded neurons (*n* = 27). **F**, Raster plot (50-ms bins) of IPSCs in a DMH neuron before, during, and after photostimulation of the SCN^NMS^→DMH input (bin duration: 50 ms). **G**, Expression of ChR2 (in red) in DMH VGat expressing neurons (scale bar: 250 µm). Photostimulation of DMH terminals in VLPO (DMH^Vgat^→VLPO) produced photo-evoked IPSCs in VLPO neurons that are blocked by bicuculline (BIC 20 µM). 30 individual IPSCs (grey) and average IPSC (black). **H**, Labelling of ChR2 (in green) in SCN NMS expressing neurons (scale bar: 200 µm) and fluorescent CTB (F-CTB, in red) in VLPO.

### Presynaptic inputs to SCN^NMS^ neurons

While previous studies using conventional tracing techniques have identified inputs to the SCN, these approaches have lacked specificity, particularly with respect to defining afferents to specific SCN cell types, including SCN^NMS^ neurons. Therefore, we aimed to identify the extra-SCN sources of presynaptic inputs to SCN^NMS^ neurons using monosynaptic conditional retrograde rabies tracing^13^. To do so, we first injected a mix of two viral vectors containing the helper constructs (EF1a-FLEX-TVA-mCherry-AAV8 and CAG-FLEX-RG-AAV8) for pseudotyped modified rabies (EnvA-DG-EGFP) into the SCN of NMSiCre mice. Four weeks later, we injected EnvA-DG-EGFP directly into the SCN (Extended Data Fig. 2a). As shown in Extended Data Fig 2b,c, the starter population of NMS neurons was restricted to the SCN. Notably, we found retrogradely labeled neurons in several brain regions, including: the lateral septum (LS), the bed nucleus of the stria terminalis (BNST), the medial preoptic (MPO), the VMPO, the anteroventral periventricular nucleus (AVPe), the paraventricular thalamic nucleus (PVA), the ventromedial hypothalamic nucleus (VMH), the arcuate nucleus (ARC), the DMH, the intergeniculate leaflet (IGL) and scattered throughout the retina (Extended Data Fig. 2d-i), the latter marking SCN^NMS^ as retinorecipient. These findings are broadly consistent with other studies describing afferents to the SCN in hamsters^14^, rats^15–17^, and mice^9^.

### SCN^NMS^ neurons are preferentially active during wakefulness

To investigate the real-time activities of SCN^NMS^ neurons across the spontaneous sleep–wake cycles of freely moving mice, we conducted *in vivo* fiber photometry to examine the state-dependent activity of SCN^NMS^neurons (see Extended Data Fig. 3). First, we unilaterally injected AAV10-EF1α-FLEX-GCaMP6s into the SCN of NMSiCre mice and implanted a photometry fiber immediately dorsal to the SCN^NMS^ cell bodies (Extended Data Fig. 3a,b). Our recordings of the SCN^NMS^ population Ca^2+^ signal revealed that SCN^NMS^ neurons are active during both wakefulness and REM sleep. Specifically, the mean GCaMP6s activity was significantly higher during wakefulness and REM sleep compared to NREM sleep, regardless of episode duration (Extended Data Fig. 3d). Additionally, SCN^NMS^ neurons responded immediately to light pulses (Extended Data Fig. 3f), indicating robust excitation of the SCN^NMS^ cell population upon light stimulation at the retina. These findings demonstrate that SCN^NMS^ neurons are highly responsive to light input in vivo, which is similar to what we and others have reported for SCN^VIP^ neurons^9,18^, but also, and unlike SCN^VIP^ neurons, that their activity is behavioral-state dependent. This finding is consistent with our chemogenetic findings in showing that the activity of SCN^NMS^ neurons subserves modulation of behavioral state.

### SCN^NMS^ neurons comprise four molecularly distinct subtypes

To investigate molecular subtypes of *Nms*+ SCN neurons, we profiled their genome-wide RNA expression by single-nuclei RNA-sequencing (snRNA-seq). We first crossed NMSiCre mice to a Cre reporter transgenic mice, H2b-TRAP (mCherry)^19^, to fluorescently label *Nms*+ cell nuclei. We then dissected SCN tissue from adult NMSiCre; H2b-mCherry mice, isolated H2b-mCherry+ single nuclei from the tissue samples and sequenced their RNA with Smart-seq2 as previously described^9,20^. Grouping the nuclei transcriptomes (∼cells) according to their expression of high-variance genes yielded four neuron clusters (Extended Data Fig. 4a). Across these cell clusters, we detected 5,855 +/- 2,756 genes per cell (mean +/- standard deviation, SD), based on 1,843,482 +/- 1,408,505 sequencing reads per cell (mean +/- SD) and essentially no contamination by mitochondrial transcripts (Extended Data Fig. 4b). The clusters likely represent neurons as they expressed neuronal marker genes (NeuN/*Rbfox3*, *Syn1*) but relatively little to no glial marker genes (oligodendrocytes/*Olig1*, polydendrocytes/*Cspg4*, macrophages/*Aif1*, astrocytes/*Agt*; Extended Data Fig. 4c). Further, cluster n01 is most likely an SCN neuron subtype, since it expresses the SCN-enriched genes *Lhx1* and *Vipr2*, but not the genes *Reln* or *Ecel1*, which are enriched in brain regions neighboring the SCN (Extended Data Fig. 4c). Differential gene expression analysis across clusters identified many cluster-enriched genes (Extended Data Fig. 4d). KEGG (Kyoto Encyclopedia of Genes and Genomes) analysis of these differentially expressed genes showed significant enrichment in cluster n01 of transcripts related to circadian rhythm (*Per3*, *Rorb*, *Rora*) and circadian enrichment (*Per3*, *Prkca*, *Nos1ap*; Extended Data Fig. 4e). These results together identify a molecularly distinct subtype of *Nms*+ neurons in the SCN which may play a role in circadian entrainment.

## Discussion

In this study, we initially demonstrate that acute activation of SCN^NMS^ neurons in vivo through chemogenetic and optogenetic methods strongly promotes wakefulness (arousal). Additionally, using fiber photometry, also in vivo, we observe that SCN^NMS^ neurons display a strong wake-active profile. Furthermore, we investigated the effect of genetically-targeted ablation of SCN^NMS^ neurons in adult NMSiCre mice, finding that it disrupted the sleep-wake cycle. Ablation specifically led to a reduction in wakefulness during the active (dark) period, especially in the WMZ, and abolished the circadian rhythm of Tb. Anterograde labeling revealed that the dorsomedial hypothalamic nucleus (DMH) was a likely key post-synaptic target of SCN^NMS^ neurons. Substantiating this, photostimulation of SCN^NMS^ terminals within the DMH robustly induced arousal from sleep. CRACM-based analysis confirmed that stimulation of SCN^NMS^ terminals within the DMH generated robust, short-latency post-synaptic inhibitory currents. In addition to these post-synaptic effects we sought to determine, using monosynaptic rabies mapping, pre-synaptic inputs to SCN^NMS^ neurons. This work revealed inputs from brain regions implicated in both the photic and non-photic regulation of the SCN clock. Unexpectedly, we also observed a strong input from the preoptic area (POA), which itself receives substantial inhibitory input from the DMH. This suggests a possible circuit basis (SCN->DMH->POA->SCN) by which SCN^NMS^ neurons can effectively promote arousal, as well as how they may become ‘silenced’ during the rest period. Finally, we conducted a detailed analysis of the transcriptional profile of SCN^NMS^ neurons using single-nuclei RNA-Seq, which revealed three distinct subtypes. Collectively, our results seem to be the first to establish a connection between molecularly-defined SCN neurons and the regulation of sleep-wake patterns, the Tb rhythm, and arousal control.

It is worth further note that the canonical two-process model posits that two distinct, interacting mechanisms form the neural basis for sleep and arousal regulation: homeostatic sleep drive (Process S) and circadian rhythms (Process C)^21–23^. While much is known about Process S, comparatively little is known about Process C. Our findings raise the intriguing possibility that SCN^NMS^ neurons represent a cellular substrate for Process C.

In summary, this comprehensive study establishes a novel connection between a molecularly-defined SCN cell population, SCN^NMS^, and the regulation of sleep-wake patterns, body temperature rhythms, and arousal control, highlighting their rapid arousal-promoting capabilities. This work therefore fills a long-standing gap in understanding of how the cellular SCN maintains optimal levels of arousal, which is the *sine qua non* for all purposive behaviors. These findings have important implications for understanding sleep disorders and circadian disruptions.

## Methods

### Animals

All animal experiments and procedures were approved and conducted following the guidelines established by the Institutional Animal Care and Use Committee of the University of California, Davis (approved protocol ID #22308) and Beth Israel Medical Center, Boston (approved protocol ID #070-2020-23). Mice were bred under controlled temperature and humidity conditions (22 ± 1 °C and 30-70% humidity) and maintained on a 12 h light/dark cycle (lights-on at 7 AM, Zeitgeber time: ZT0). Food and water were available ad libitum throughout the study.

Male heterozygous *NMS-IRES-Cre* mice at 8-12 weeks of age were used for behavioral experiments. NMS-IRES-Cre mice (sourced from Jackson Laboratory: C57BL/6-Tg (Nms-icre)20Ywa/J, Cat. #027205) express codon-improved Cre recombinase in a subset of Nms-expressing neurons within the suprachiasmatic nucleus^11^. Littermates mice that did not express cre were used as controls. Adult male and female *NMS-IRES-Cre* mice (6-12 weeks of age) were used for electrophysiology and anatomical studies.

### Stereotaxic brain injections

Mice were anesthetized with ketamine/xylazine (100 and 10 mg/kg, respectively, i.p.), received slow-release meloxicam subcutaneously (4 mg/kg) for pain relief. The surgical region (top of the skull) was shaved in preparation for surgery. The mouse was then secured into a stereotaxic frame and the skin over the top of the skull sterilized with betadine and 70% isopropyl ethanol before making an incision along the midline of the skull, exposing lambda and bregma. Burr holes (0.7 mm diameter) were drilled immediately through the skull above the SCN and an incision was made in the meningeal layer with a 25G needle. Subsequently, a pulled glass micropipette (10–20 μm diameter tip) containing the viral vector was slowly lowered down to the SCN [anteroposterior (AP): - 0.2 mm from bregma; lateral (ML): ± 0.2 mm; dorsoventral (DV): -5.0 mm; as per the mouse atlas of Paxinos and Franklin]^24^. The virus was injected by an air pressure system using picoliter air puffs through a solenoid valve (Clippard CR-EV-3-24-L) pulsed by a Grass S44 stimulator to control injection speed (30 nl/min). Following the intracranial injection, we waited for 3 minutes to allow dispersion of the viral vector before gently withdrawing the glass pipette from the brain. The scalp wound was closed with surgical sutures, 0.5 mL sterile saline was administered subcutaneously, and the mouse allowed to resume consciousness on a heating pad. To chemogenetic activation of the SCN^NMS^ neurons, AAV10-hSyn -DIO-hM3Dq-mCherry (80 nL per side, packaged as described previously) were bilaterally injected into the SCN of the *NMS-IRES-Cre* mice (n =7). To selective ablation of the SCN^NMS^ neurons, *NMS-IRES-Cre* mice (n = 5) and wild-type mice (WT; n =7) received bilateral injections of AAV10-CMV-FLEX-mCherry-DTA (120 nL per side) into the SCN. The DTA vector, which was created by Drs. Patrick M. Fuller and Drs. Michael Lazarus, has undergone extensive validation, and is well published ^9,25,26^. Following injections of DTA-AAV, and per prior validation studies with this viral-based toxin, we allowed at least 4 weeks for the lesion to develop before starting the recordings. To optogenetic activation of the SCN^NMS^ neurons, AAV10-EF1a-FLEX-hChR2(H134R)-eYFP (ChR2-eYFP, injection volume: 75 nL) was bilaterally injected into the SCN of the *NMS-IRES-Cre* mice. For photometry experiments, AAV10-EF1α-FLEX-GCaMP6s (40 nL) was injected into the left SCN in NMS-IRES-Cre mice.

### EEG/EMG and optical fiber implants

All mice used in behavioral experiments underwent a surgery to implant a headstage for recording the electroencephalogram (EEG) and electromyogram (EMG) and an optical fiber for optogenetic stimulation or photometry recordings. The headstage consisted of a six-pin connector (Heilind Electronics, catalog #MMX853-43-006-10-001000) soldered to four EEG screws (Pinnacle Technology Inc., catalog #8403) and two flexible EMG wire electrodes (Plastics One, catalog #E363/76/SPC). Mice were prepared for surgery as described above and burr holes were drilled immediately above the SCN (AP: -0.2 mm; ML: 0 mm) or DMH (AP: -1.55 mm; ML: 0 mm for placement of the optical fibers. 3 additional burr holes (0.7 mm diameter) were drilled in the skull: one anterior burr hole was drilled at AP: +1.0 mm and ML: - 1.0 mm and 2 posterior burr holes were drilled at AP: - 2.0 and ML: ± 1.0. The EEG electrode screws were inserted into the burr holes and each optical fiber was stereotaxically guided into position above the SCN (DV: -4.7 mm) and the DMH (DV: - 4.5 mm) and glued into place using a mixture of dental cement and cyanoacrylate glue. The EMG electrodes were guided down the back of the neck underneath the trapezius muscle. An additional layer of dental cement was then applied to the whole assembly to insulate the headset and provide structural stability. The scalp wound was closed with surgical sutures and the mouse was kept in a warm environment until resuming normal activity.

The optical fibers were manufactured in-house, comprised a fiber optic cable (200 μm outer diameter, ThorLabs FT200EMT for optogenetic experiments; 400 μm outer diameter, ThorLabs FP400URT for photometry experiments) inserted into a ferrule (ceramic, 200 μm internal diameter, ThorLabs CFLC230-10 for optogenetic experiments; stainless steel, 400 μm internal diameter, ThorLabs SFLC440-10 for photometry experiments) and epoxied into place (PFP 353ND, Precision Fiber Products). At the mating end of the ferrule, the optical fiber was flat cleaved and polished and at the other end, the optical fiber was cut to size (5.5 mm).

### Core body temperature logger implants

The core body temperature of the mice was recorded using DST nano-T temperature loggers (length 17 mm, diameter 6 mm, weight 1g; Star-Oddi, Gardabaer, Iceland). Following the headstage implantation, an incision was made in the IP cavity and the loggers were implanted into the peritoneal cavity of each mouse. The incision was closed with surgical sutures. The loggers were programmed to record body temperature at 5-min intervals and remained in place for eight weeks. Prior to implantation, and following removal from mice, all the loggers were placed in Cetylcide-G (CETYLITE) overnight for sterilization then washed with saline before implant. Mice were recorded for at least 21 days under an LD cycle, followed by at least 10 days in DD, and then returned to LD conditions for at least 7 days.

### Sleep-wake monitoring

Following a minimum of 10 days for post-surgical recovery, the mice were housed individually in transparent barrels in an insulated sound-proofed recording chamber maintained at an ambient temperature of 22 ± 1 °C and on a 12 h light/dark cycle (lights-on at 7 AM, Zeitgeber time: ZT0) with food and water available ad libitum. Mice were allowed to be habituated to the recording chambers, EEG/EMG recording cable and optical patch cables for at least 3 days before starting polygraphic recordings. Cortical EEG (bipolar, fronto-parietal, ipsilateral) and EMG signals were amplified x5000 (Model 3600; AM Systems, Sequim, WA) and digitized with a resolution of 500 Hz using a Micro 1401-3 (Cambridge Electronic Design, Cambridge, UK).

### Chemogenetic experiments

For chemogenetic experiments, mice were recorded for a 24 h baseline periods, followed by IP injections of saline (vehicle injection) or clozapine-N-oxide (CNO; NIMH Chemical Synthesis and Drug Supply Program; 0.3 mg/kg or 1 mg/kg in saline). Injections were performed at 11:00 A.M. (ZT4; light period, time of high sleeping drive), in a randomized cross-over design, with each injection separated by a 4-day washout period.

### Optogenetic experiments

For optogenetic experiments, experiments were carried out between ZT3-ZT9 (a period of low activity in nocturnal mice). We modified an open-source Online Sleep Detection script (Spike 2, Cambridge Electronic Design, source code available from the Cambridge Electronic Design website) to detect when mice had been in NREM sleep or REM sleep wake for at least 20 s, to trigger a 10 s duration, 12-15 mW, 473 nm blue light (R471003GX, LaserGlow, Toronto, Canada) stimulation to the SCN or DMH. The stimulation consisted of light pulses (5 ms) delivered at a variety of frequencies (1 Hz, 5 Hz, 10 Hz, 20 Hz or a sustained 2 s stimulation), delivered in a random order throughout the recording session. A refractory period of at least 3 minutes was allowed between each stimulation, no mouse received more than 200 stimulations over any recording session and each recording session was separated by at least two days. Simulations triggered in NREM versus REM were carried out in separate recording sessions.

### In vivo fiber photometry

To simultaneously record the GCaMP6s signal, light from a 465 nm blue LED (the excitation wavelength for GCaMP6s, LEDC1-B_FC Doric) and a 405 nm UV LED (LEDC1-405_FC) was directed to a fiber optic mini-cube containing UV and GFP filters (FMC5_AE(405)_ AF(420-450)_E1(460-490)_F1(500-550)_S, Doric) via a 200 μm, 0.22 NA fiber patch cable. A sample patch cord (400 μm, 0.48 nm) was mated to the photometry fiber ferrule on the mouse’s head to deliver blue and UV light to the SCN and collect the emitted fluorescence from GCaMP6s-expressing cells. This emitted signal was directed back through the filter cube to two photodetectors (Newport 2151) via 600 μm, 0.66 NA fiber patch cables. The signal was amplified at the photodetector (DC gain: 2 x 10^10^) and digitized at 500 Hz (Micro-1401-3, Cambridge Electronic Design). Recordings in which UV excitation of the tissue resulted in a moving signal that ran parallel to the recorded calcium dependent GCaMP6s signal were assumed to be contaminated with movement artifact and discarded from the analysis.

For simultaneous EEG/EMG recordings, the EEG/EMG headset was connected, via a low weight custom-made cable, to a freely moving electrical commutator. Following habituation, mice were recorded for a 6-hour period between ZT 3-9.

### Sleep scoring and analysis

Sleep scoring was carried out using Spike 2 (CED). For optogenetic experiments, polysomnographic records were visually scored in 5 s epochs for wakefulness (W), rapid eye movement (REM) and non-REM (NREM) sleep. The latency to wake and length of wake episodes following the optogenetic stimulation were calculated and exported from Spike for visualization using a custom written Python script (https://www.python.org)^12^.

For photometry experiments, an investigator that was blinded to the GCaMP6s signal marked the transition points between each sleep-wake state. Sleep-wake states were identified using a combination of observed EEG delta power, theta power, theta: delta ratio and EMG power. At wake to NREM transitions, the initiation of NREM was identified by a 5 s period of continuous high power delta activity. At NREM to REM transitions, REM was identified by the clear onset of theta activity, together with a cessation of high-power delta activity. After behavioral sleep stage scoring, the GCaMP6s signal and sleep staging timestamps were exported to MATLAB for further analysis. The data was down sampled to 1 Hz and fitted to a second order exponential curve to correct for bleaching. This curve was designated as F_0_ and used to calculate ΔF/F = F_t_-F_0_/F_0_, where F_t_ denotes the GCaMP6s signal at time t. To normalize fluorescence levels between mice, ΔF/F was then processed as a z-score. At transitions, the GCaMP6s signal was zeroed by calculating the median value of the 60 s preceding the transition and subtracting this from the GCaMP6s signal throughout the transition (i.e., 60 s before to 60 s after the transition occurred). To quantify the effect of the state transition upon GCaMP6s activity, the GCaMP6s signal was averaged 10 s immediately preceding the state transition and compared against the GCaMP6s signal averaged 10 s after the state transition. To quantify GCaMP6s activity during different arousal states, the GCaMP6s signal was averaged over the duration of all wake, NREM or REM sleep episodes.

### Rabies tracing

For monosynaptic retrograde modified rabies tracing we first made injections of a mixture of 60 nL AAV8-EF1ɑ -DIO-TVA and 60 nL AAV8-CAG-DIO-RG into the SCN as described above (Stereotaxic brain injections). Four weeks after the initial surgeries, 200 nL of EnvA-ΔG-eGFP was injected into the same place. Mice were perfused 11 days following the second stereotaxic injection and processed for histology (as described in Immunohistochemistry). In brief, one entire series was immunolabelled for dsRed to allow visualization of TVA-mCherry and then mounted. Neurons containing both TVA-mCherry and eGFP were considered the starter population. Each eGFP positive neuron in the brain was counted and registered to the mouse atlas of Paxinos and Franklin^24^. The profile of afferent inputs is presented as the number of eGFP expressing neurons within a particular brain region as a percent of the total number of eGFP neurons counted in the whole brain (the input fraction).

### Data analysis for body temperature

Data retrieval from the temperature loggers was performed using Mercury v. 6.14 software and the associated Communication Box (Star-Oddi, Gardabaer, Iceland). Tb data were first analyzed and plotted using Clocklab (Coulbourne Instruments, Natick, MA). For each mouse, we obtained Tb rhythm period (Chi-squared periodogram) for the last 10 days in LD and the last 10 days in DD. Double plots of the body temperature during the constant darkness were created by Python as previously described with some modifications^27^. Briefly, the body temperature was normalized and fitted to the grayscale. In each double plot, the lowest body temperature of each subject was colored in white, and 93% or higher of the maximum body temperature was colored in black. Each bin indicated 5 minutes. The acrophase of the Tb rhythms were calculated from a least squared fit during the last 10 days of LD. Chi-square periodograms were generated from the last 10 days data in constant darkness. The largest peak in the periodogram in the range of 12–36 h was selected as the circadian period.

### In vitro CRACM recording

To study the SCN^NMS^ → DMH → VLPO circuit, AAV10-EF1a-DIO-hChR2(H134R)-eYFP (ChR2-eYFP, injection volume: 100 nL) was injected bilaterally into the SCN of *NMS-IRES-Cre* mice (n = 5 mice). In these mice we also placed a second microinjection of Alexa Fluor-555-conjugated Cholera Toxin Subunit B (F-CTB; Invitrogen) into the ipsilateral VLPO (75 nL; AP: 0.36 mm, DV: -5.0 mm, ML: 0.70 mm) to retrogradely label DMH neurons projecting to the VLPO. In a second set of experiments, ChR2-eYFP (60 nL) was injected unilaterally into the DMH (AP: -1.82 mm; ML:0.3 mm; DV:-5 mm) of VGat-IRES-Cre mice. Five to six weeks after ChR2-eYFP and F-CTB injections, mice were sacrificed for *in vitro* electrophysiological recordings of DMH or VLPO neurons. Mice were deeply anaesthetized with isoflurane (5% in oxygen) via inhalation and transcardially perfused with ice-cold cutting ACSF (N-methyl-D-glucamine, NMDG-based solution). The mouse brains were then quickly removed, and 250 μm-thickness coronal slices were made in ice-cold NMDG-based ACSF using a vibrating microtome (VT1200S, Leica). We first incubated the slices containing the DMH or VLPO for 5 min at 37°C, then transferred them into a holding chamber at 37°C containing normal ACSF (Na-based solution) for 10 minutes. Following this two-step incubation period, we let the brain slices gradually return to room temperature (∼ 1 hour) before transferring them to the recording chamber where they were submerged and perfused (1.5 ml/min) with Na-based ACSF. Recordings were conducted under infrared differential interference contrast (IR-DIC) visualization and we recorded F-CTB-containing DMH neurons using a combination of fluorescence and IR-DIC microscopy. We used a fixed stage upright microscope (BX51WI, Olympus America) equipped with a Nomarski water immersion lens (Olympus 40X / 0.8 NAW) and IR-sensitive CCD camera (ORCA-ER, Hamamatsu, *Bridgewater, NJ*). Real-time images were acquired using MATLAB (MathWorks) script software. We recorded DMH and VLPO neurons in whole-cell configurations using a Multiclamp 700B amplifier (Molecular Devices, Foster City, CA), a Digidata 1322 A interface, and Clampex 9.0 software (Molecular Devices). Neurons showing a greater than 10% change in input resistance over the duration of the recording were excluded from the analysis. For all recordings, we recorded at a holding potential of 0mV and using a Cs-methane-sulfonate-based pipette solution. We photostimulated axons terminals expressing ChR2 in the DMH and VLPO using full-field light pulses (∼10 mW/mm^2^, 1 mm beam width) from a 5W LUXEON blue light-emitting diode (470 nm wavelength; #M470L2-C4; Thorlabs), coupled to the epifluorescence pathway of the microscope. To record IPSCs, we applied 10 ms light pulses (0.1 Hz, for a minimum of 30 trials) to the brain slice in order to elicit photo-evoked IPSCs. At the end of the recordings, recorded slices and adjacent slices containing ChR2 expressing neurons and F-CTB injection sites were fixed overnight in 10% buffered formalin. The location of the injection sites was verified using a Leca Stellar 5 confocal microscope.

### Solutions for CRACM experiments

ACSF NMDG-based solution contained (in mM): 100 NMDG, 2.5 KCl, 1.24 NaH_2_PO_4_, 30 NaHCO_3_, 25 glucose, 20 HEPES, 2 thiourea, 5 Na-L-ascorbate, 3 Na-pyruvate, 0.5 CaCl_2_, 10 MgSO_4_ (pH 7.3 when carbogenated with 95% O_2_ and 5% CO_2_; 310-320 mOsm). ACSF Na-based solution contained (in mM): 120 NaCl, 2.5 KCl, 1.3 MgCl_2_, 10 glucose, 26 NaHCO_3_, 1.24 NaH_2_PO_4_, 4 CaCl_2_, 2 thiourea, 1 Na-L-ascorbate, 3 Na-pyruvate (pH 7.3-7.4 when carbogenated with 95% O_2_ and 5% CO_2_; 310-320 mOsm). The Cs-methane-sulfonate-based pipette solution contained (in mM): 125 Cs-methane-sulfonate, 11 KCl, 10 HEPES, 0.1 CaCl_2_, 1 EGTA, 5 Mg-ATP and 0.3 Na-GTP (pH adjusted to 7.2 with CsOH, 280 mOsm).

### Data analysis for electrophysiological recording

Recording data were analyzed using Clampfit 10 (Molecular Devices), MiniAnalysis 6 (Synaptosoft, Leonia, NJ) and MATLAB (MathWorks) software. We prepared the figures using Igor Pro version 6 (WaveMetrics), Prism 7 (GraphPad, La Jolla, CA) and Photoshop (Adobe) software. To ensure unbiased detection of the synaptic events, the IPSCs were detected and analyzed automatically using MiniAnalysis. We considered DMH and VLPO neurons to be responsive to photostimulation if their IPSC probability during the first 50 ms that follows the light pulses was greater than the IPSC probability + five times the SEM in baseline conditions (baseline IPSC probability = 7.08 ± 1.32%, n = 27)^28^. We calculated the latency of the photo evoked IPSCs as the time difference between the start of the light pulse and the 5%rise point of the first IPSC^29^.

### Immunohistochemistry

At the end of experiments, mice were sacrificed by deep anesthesia with isoflurane (5% in oxygen) via inhalation, and transcardially perfused with 30 mL saline, followed by 40 mL of neutral phosphate buffered formalin (4%, Fischer Scientific). Brains were removed from the skull and incubated in neutral phosphate buffered formalin for 2-hr, followed by 20% sucrose at 4°C until they sank, and then sectioned at 40 μm on a freezing microtome into three series. Sections were washed in phosphate buffered saline (PBS), and incubated in primary antiserum (mCherry rat monoclonal antibody (1:1K; Invitrogen; Catalog #M11217;), c-Fos rabbit polyclonal antibody (1:3K; Millipore; catalog #ABE457), GFP chicken polyclonal antibody (1:1K; Invitrogen; catalog #A10262) diluted in PBS containing 0.3% Triton X-100 (PBST) and 0.2% sodium azide for overnight at room temperature. Sections were washed 3 times in PBS and incubated in fluorescent secondary antiserum for 2 hours. For all secondary antibody immunohistochemical controls, the primary antibodies were omitted and the tissue showed no immunoreactivity above background. Secondary antibodies included the following: Donkey anti-rabbit Alexa Fluor 488 (1:1000; Invitrogen); Donkey anti-rat Alexa Fluor 594 (1:1000; Invitrogen); Goat anti-rabbit Alexa Fluor 568 (1:1000; Invitrogen); and Goat anti-chicken Alexa Fluor 488 (1:1000; Invitrogen). Finally, the sections were washed twice in PBS before being mounted on positive charged slides (Denville Scientific, Inc) that had been pre-subbed in 1% gelatin solution, counter slipped using VECTASHIELD Vibrance® Antifade Mounting Medium with DAPI (Vector Laboratories; catalog #H-1800-10) and visualized with a florescence microscope (Keyence BZ-X710, Japan).

### Image analysis

For retrograde tracing with EnvA-DG-EGFP, slides were visualized on OlyVIA software and EGFP-expressing neurons from every section of one series were counted manually and registered according to the atlas of Franklin and Paxinos^24^. The profile of afferent input is presented as the number of EGFP expressing neurons within a particular brain region as a percent of the total number of EGFP neurons counted in the brain (the input fraction). The input fraction for each brain area was averaged over all cases.

### Single cell RNA sequencing data analysis

sNuc-Seq Data Generation: Six-week old (juvenile; n = 3) and 10-week old (adult, n = 3) NMS-IRES-Cre::H2b-TRAP male siblings were bred from the same NMS-IRES-Cre x H2b-TRAP mouse line. One hour after light cycle onset, the NMS-IRES-Cre::H2b-TRAP mice were removed from their home cage and rapidly decapitated in order to minimize stress-related changes in gene expression. Brains were extracted quickly, chilled briefly in ice-cold DMEM/F12 media, and then sectioned coronally at 1-mm intervals in a chilled stainless steel brain matrix. The brain sections containing the SCN and sections adjacent to the SCN were collected in ice-cold RNAlater (Qiagen catalog # 76106). Each section was imaged using a fluorescence stereoscope (Zeiss Discovery V8) to visualize fluorescently labeled cells in the SCN for dissection. Dissected tissue was pooled across mice by age group (juvenile, adult) and placed into ice-cold RNAlater overnight in the dark at 4 °C. The following morning, the RNAlater-preserved tissue samples were processed into single-nuclei suspensions according to the sNuc-Seq method of Habib et al.^20^. In brief, the two SCN tissue samples (juvenile, adult) were Dounce homogenized (Wheaton tissue grinders, catalog # 357538) and separated by density gradient centrifugation as described in the published protocol^20^. Each pellet was then re-suspended in 0.5 mL sNuc-Seq resuspension buffer and filtered through a 20 μm mesh (Miltenyi Biotec pre-separation filters, catalog # 130-101-812) to eliminate clumped nuclei and larger debris. The nuclei were counterstained with two drops of NucBlue Live ReadyProbes Reagent (ThermoFisher Scientific catalog # R37605) per mL of nuclei. Nuclei were sorted by MoFlo Astrios EQ cell sorter (Harvard Bauer core facility), gating for NucBlue + /mCherry+ singlets and then individually distributing them into wells of 96-well plates containing 5 μL per well of nuclei capture buffer (1% beta-mercaptoethanol in Qiagen Buffer TCL; catalog # 1031576). Each plate was loaded with equal numbers of samples from juvenile and adult SCN tissue. Subsequently, the loaded plates were centrifuged for 5 min at 2500 Å∼ g and 4 °C to ensure the nuclei effectively reached the capture buffer and were then promptly frozen at −80 °C for up to 4 weeks prior to utilization. The sample plates were transferred to an RNA-dedicated PCR cabinet, where they were rapidly thawed, and immediately purified using RNA-Clean XP (Beckman Coulter, catalog # A63987) at a ratio of 2.5:1 of RNA-Clean XP to the sample volume. The elution process involved using 4 μL of Smart-seq2 buffer (consisted of 2 μL Smart-seq2 cell lysis buffer, 1 μL 100 μM oligo-dT primer, and 1 μL 10mM dNTP mix). The complementary DNA (cDNA) was then generated and amplified according to the Smart-Seq2 protocol ^30^, except that 22 PCR cycles were used during the amplification step. After the PCR step, 10 μL of each sample was transferred to fresh 96-well plates for subsequent processing. The amplified cDNA was purified with two cycles of SPRIselect (Beckman Coulter, catalog # B23318; at a ratio of 0.8:1 SPRIselect to sample volume), with elution performed in ultrapure water. cDNA concentration was measured by Qubit fluorometry in a microplate configuration, where each well contains 2 μL of cDNA sample and 98 μL of Qubit working solution (ThermoFisher Scientific, catalog # Q32854). Qubit readings were obtained with a fluorescence microplate reader (Molecular Devices SpectraMax M5 at the Harvard Medical School ICCB-Longwood core facility) and used to calculate DNA concentrations by comparing each sample’s Qubit reading to standards in a 50% dilution series (ranging from 0.25–0.01 ng/μL alongside a negative control, 0.0 ng/μL). The cDNA samples were subsequently diluted to 0.15 ng/μL with ultrapure water and processed into sequencing libraries with sample-specific indices using the Nextera DNA Sample Prep Kit and Nextera XT Index Kit v2 sets A–D (Illumina, catalog #s FC-131-1096, FC-131-2001, FC-131-2002, FC-131-2003, FC-131-2004, respectively). The Nextera XT was used at 1/4 reaction volumes volumes while adhering to the guidelines provided by manufacturer. Sequencing libraries were purified with two rounds of SPRIselect (0.8:1 ratio of SPRIselect to sample volume) and eluted in a single volume of ultrapure water. The concentration of the library pool concentration was determined usin Qubit as per the manufacturer’s instructions, assessed for size using Agilent BioAnalyzer high sensitivity kit, and then diluted with ultrapure water to achieve a size-corrected concentration of 2 nM. The diluted library pool was denatured and was further diluted to 2pM loading concentration in accordance with Illumina NextSeq recommendations, then sequenced with a NextSeq 500 v2 75cycle sequencing kit (Illumina, catalog # FC-404-2005).

sNuc-Seq data processing. Base calls and demultiplexing were performed with bcl2fastq2 v2.20.0. Reads were aligned to the mouse genome (build: mm10, GRCm38) by HISAT2 v2.1.0 (using “-k 2”)^31^. Aligned reads were converted to BAM using SAMtools v1.8.0^32^. Duplicate and low-quality reads were removed with Picard 2.17.0 (http://broadinstitute.github.io/picard). Reads were assigned to features (genes) using the feature Counts from the Subread package v1.6.0 (using recommended setting and a custom meta-annotation that merges overlapping isoforms)^33^. Ultimately, expression values were calculated by counting the reads assigned to each gene.

sNuc-Seq data analysis. Clustering and cluster marker analyses were performed in Seurat v2.3.4^34^ using the standard workflow and default settings, unless specified otherwise. First, the gene expression values were imported and log normalized. Highly variable genes were identified with Seurat’s FindVariableGenes function based on their higher ratios of dispersion in relation to mean expression. These identified highly variable genes were used for principal component analysis (PCA) with Seurat’s RunPCA function. The transcriptomes were then clustered across the first 10 principal components (PCs) using Seurat’s FindClusters function, with a resolution of 1. The clusters were numbered based on their position in hierarchical clustering dendrogram. To identify cluster markers, differential expression analysis was performed via Seurat’s FindAllMarkers function with default settings. Clusters were categorized as “SCN” or “non-SCN” based on the anatomical expression patterns of top cluster markers as observed in the Allen Mouse Brain Atlas^35^ (Lhx1; Vipr2; Reln; Ecel1). The SCN clusters were re-clustered by excluding the non-SCN cluster and then repeating the workflow described above, beginning with the variable gene selection. Plots were generated with Seurat v2.3.4 and pheatmap v1.0.12 packages for R.

### Statistics

All statistical analysis were performed using Prism v8 (GraphPad Software, San Diego, CA, United States) and details of the statistical tests used for each experiment may be found in the figure legends. In all cases, values showing p < 0.05 were considered significant. Sample size and power calculations were performed *post hoc* using PS Power and Sample Size Calculations by William D. Dupont and Walton D. Plummer (http://biostat. mc.vanderbilt.edu/wiki/Main/PowerSampleSize) using means and standard deviations derived from our analysis. All experimental data were subject to histological validation. Data were excluded if the conditions of the histological validation were not met (i.e., cases in which transduction of the viral vector extended beyond the boundaries of the SCN or optical fiber placement was not correctly positioned). Sample sizes reported in the manuscript reflect only data that was confirmed by histological validation. All behavioral recordings were scored by an investigator that was blinded to the recording conditions.

### Reporting summary

Further information on research design is available in the Nature Portfolio Reporting Summary linked to this article.

## Data availability

All data needed to evaluate the conclusions in the paper are available in the main text, figures and extended data.

## Acknowledgments

We thank Quan Hue Ha, Emily T Doisy, Ebun Smith for superb technical assistance. We thank Drs. Christelle Anaclet and Heinrich Gompf for helpful discussions. We also thank Dr. Fuyuki Asano for body temperature data analysis using Python. This work was supported by National Institutes of Health grants R01NS118856 and R01NS073613 (to P.M.F.) and R01NS091126 (to E.A.).

## Author contributions

Conceptualization, Y.E.W., R.D.L., E.A., and P.M.F.; Methodology, Y.E.W., R.D.L., R.Y.B., A.V., S.S.B., J.C. and E.A.; Investigation, Y.E.W., R.D.L., A.V., L.T.S., S.S.B., E.A.; Writing – Original Draft, Y.E.W., and P.M.F.; Writing – Review & Editing Preparation, Y.E.W., R.D.L., R.Y.B., A.V., L.T.S., S.S.B., D.S., J.C., E.A., and P.M.F.; Visualization, Y.E.W., R.D.L., A.V.; Supervision, E.A., and P.M.F.; Project Administration, Y.E.W., R.D.L., E.A., and P.M.F.; Funding Acquisition, E.A., and P.M.F.

---

**Extended Data Fig. 1.**
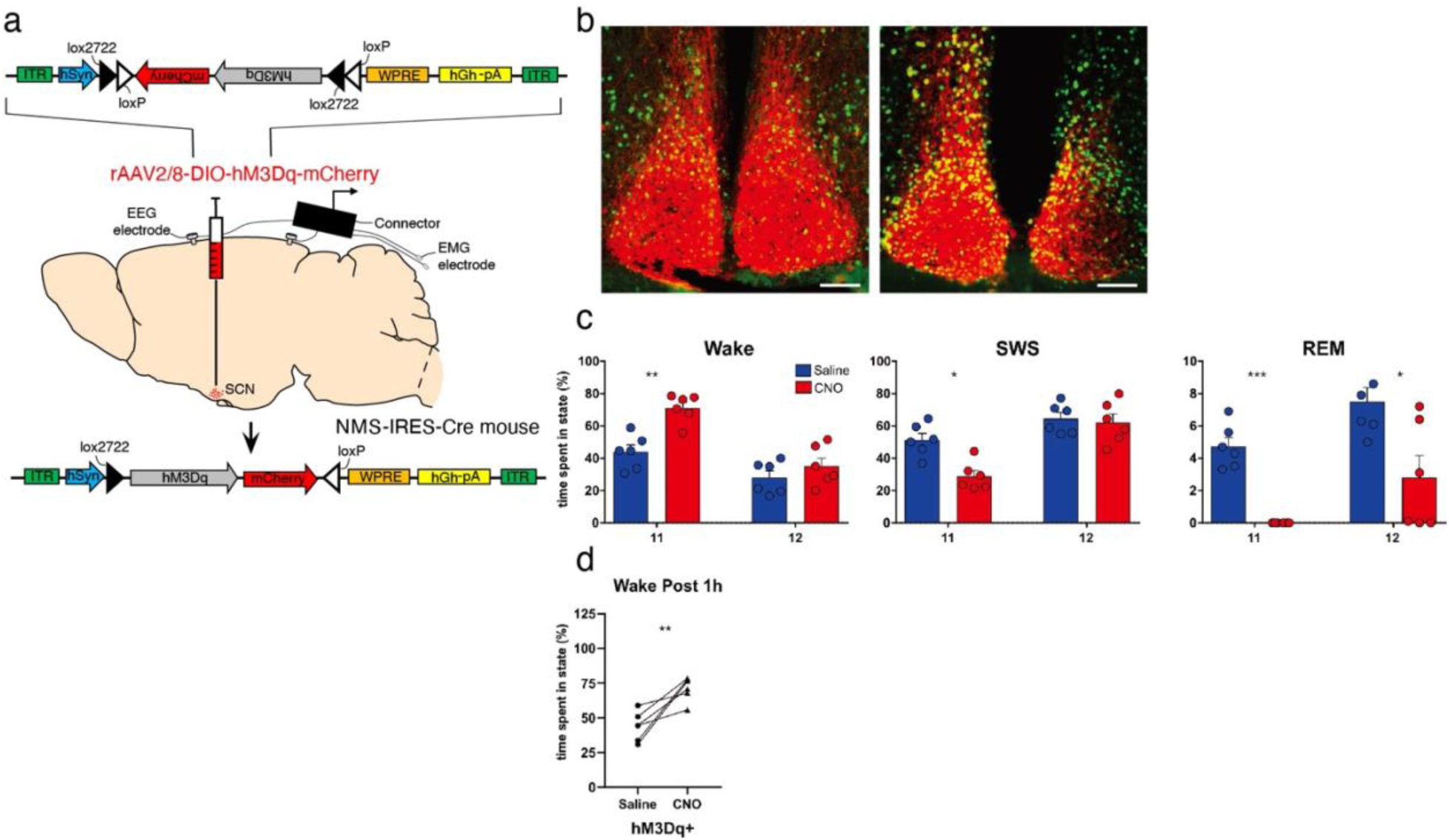
Chemogenetic activation of SCN^NMS^ neurons increases wakefulness. **a**, Schematic illustration of the experimental setup: AAV-hSyn-DIO-hM3Dq-mCherry was bilaterally injected into the SCN of NMS-IRES-Cre mice. Two weeks later, the mice were equipped with an EEG/EMG headset to record sleep-wake patterns. **b**, Photomicrograph depicting immunofluorescent expression of hM3Dq-mCherry (Red) and cFos (green) in SCN^NMS^ neurons. Scale bar, 200μm. Clear co-localization of mCherry expression and cFos induction, indicating activation of hM3Dq-expressing neurons in response to CNO. **c**, Total time spent in wakefulness, NREM sleep, and REM sleep during the 1^st^ h and 2^nd^ h recording following the saline or CNO administration (n = 6 mice). Two-way ANOVA followed by Sidak post hoc test is shown. *p < 0.05; **p < 0.01; ***p < 0.001. **d**, Total time spent in wakefulness during the 1^st^ h recording following the saline or CNO administration (n = 6 mice). Values are the mean ± SEM for each group (n = 6 mice). Two-way ANOVA followed by Sidak post hoc test is shown. **p < 0.01.

**Extended Data Fig. 2.**
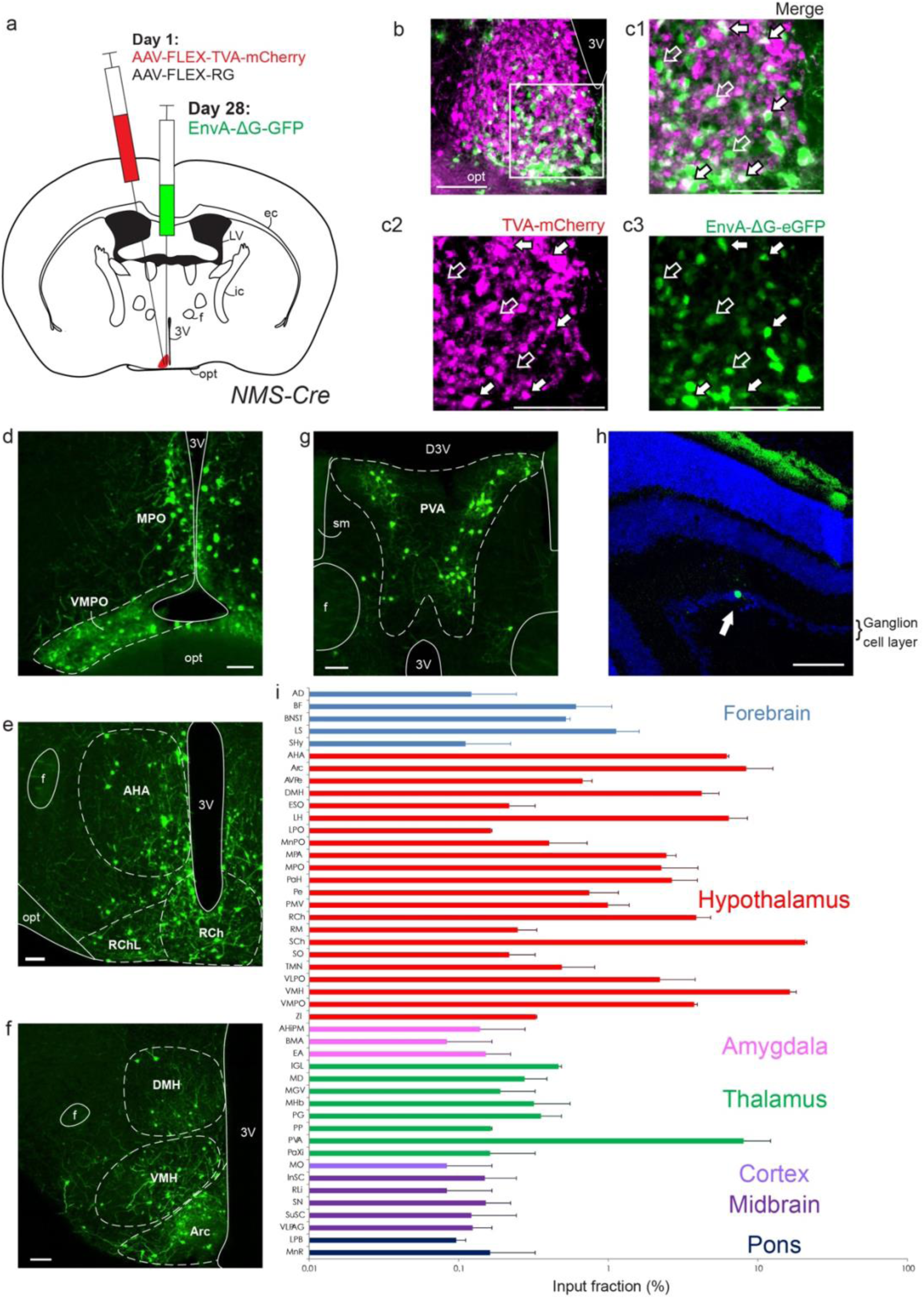
Distribution of inputs to SCN^NMS^ neurons. **a**, Schematic of unilateral stereotaxic injections of the two starter viruses (AAV-FLEX-TVA-mCherry and AAV-FLEX-RG) into the SCN, then 3 weeks later unilateral injections of pseudotyped modified rabies (EnvA-ΔG-eGFP) into the same location. **b,c**, Photomicrographs showing injection site 3 weeks after rabies injection. TVA-mCherry transfected neurons are labeled in magenta. Starter population neurons, i.e. neurons with both TVA-mCherry and EnvA-ΔG-eGFP, are labeled in white. SCN input neurons expressing EnvA-ΔG-eGFP are labeled in green. C1-C3 show boxed area in B. Filled arrows point towards starter population neurons, while unfilled arrows point towards neurons expressing only EnvA-ΔG-eGFP or TVA-mCherry. Scale bar: 100µm. **d,e,g**, Photomicrographs showing EnvA-ΔG-eGFP expressing input neurons to the SCN in the medial preoptic nucleus and ventromedial preoptic nucleus (d), anterior hypothalamic area and retrochiasmatic area (e), dorsomedial hypothalamic nucleus, ventromedial hypothalamic nucleus, and arcuate hypothalamic nucleus (f), and paraventricular thalamic nucleus. Scale bar: 100µm. **h**, Photomicrograph showing EnvA-ΔG-eGFP expressing input neuron (green, filled arrow) in the ganglion cell layer of the retina, counter-stained with DAPI. Scale bar: 100µm. **i**, Bar chart depicting the number of inputs to the SCN as a percentage of the total number of input neurons (n=2 cases, error bars indicate mean ± S.E.M. x-axis shown as a logarithmic scale). Abbreviations: 3V; 3rd ventricle, AD; anterodorsal thalamic nucleus, AHA; anterior hypothalamic area, AHiPM; amygdalohippocampal area, posteromedial part, Arc; arcuate hypothalamic nucleus, AVPe; anteroventral periventricular nucleus, BF; basal forebrain, BMA; basomedial amygdaloid nucleus, BNST; bed nucleus of the stria terminalis, D3V; dorsal 3^rd^ ventricle, DMH; dorsomedial hypothalamic nucleus, EA; extended amygdala, ec; external capsule, ESO; episupraoptic nucleus, f; fornix, ic; internal capsule, IGL; intergeniculate leaflet, InSC; intermediate stratum, LH; lateral hypothalamic area, LPB; lateral parabrachial nucleus, LPO; lateral preoptic area, LS; lateral septal nucleus, LV; lateral ventricle, MD; mediodorsal thalamic nucleus, MGV; medial geniculate nucleus, ventral part, MHb; medial habenular nucleus, MnPO; median preoptic nucleus, MnR; median raphe nucleus, MO; medial orbital complex, MPA; medial preoptic area, MPO; medial preoptic nucleus, opt; optic tract, PaH; paraventricular hypothalamic nucleus, PaXi; paraxiphoid nucleus of thalamus, Pe; periventricular hypothalamic nucleus, PG; pregeniculate area, PMV; premammillary nucleus, ventral part, PP; peripeduncular nucleus, PVA; paraventricular thalamic nucleus, RCh; retrochiasmatic area, RChL; retrochiasmatic area, lateral part, RLi; rostral linear nucleus, RM; retromammillary nucleus, SCh; suprachiasmatic nucleus, SHy; septohypothalamic nucleus, sm; stria medullaris, SN; substantia nigra, SO; supraoptic nucleus, SuSC; superficial stratum of the superior colliculus, TMN; tuberomammillary nucleus, VLPAG; ventrolateral periaqueductal gray, VLPO; ventrolateral preoptic nucleus, VMH; ventromedial hypothalamic nucleus, VMPO; ventromedial preoptic nucleus, ZI; zona incerta.

**Extended Data Fig. 3.**
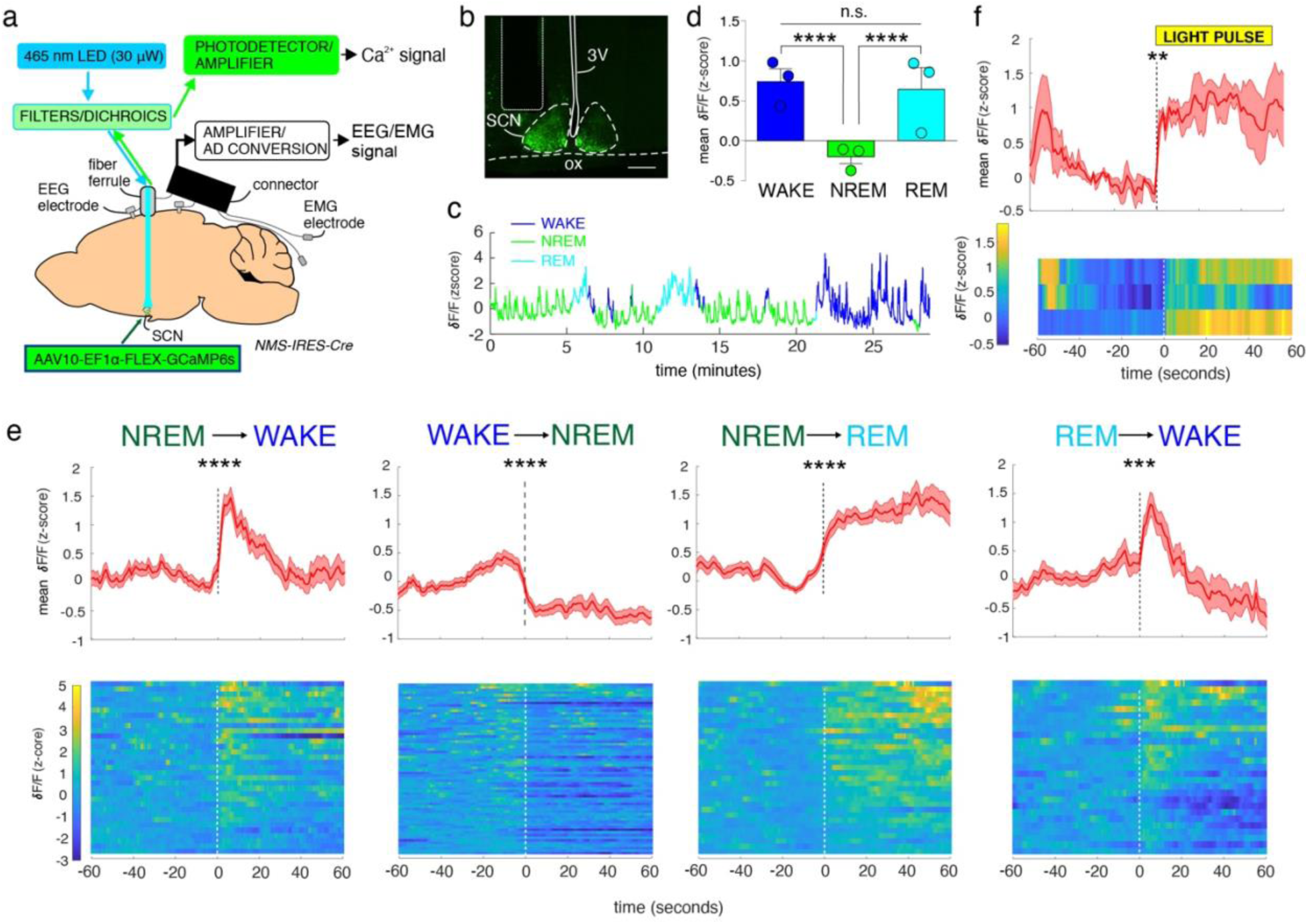
In vivo fiber photometry of SCN^NMS^ neurons during natural sleep-wake behavior and during a light pulse. **a**, Experimental schema depicting viral vector injection of AAV10-EF1α-FLEX-GCaMP6s into the SCN of a *NMS-IRES-Cre* mouse and surgically implanted photometry fiber for recording population Ca^2+^ activity and EEG/EMG head stage for recording sleep-wake. **b**, Representative photomicrograph of GCaMP6s containing neurons (green) and the tip of the photometry fiber in the SCN of a *NMS-IRES-Cre* mouse. Scale bar: 200 µm. ox; optic chiasm, 3V; 3rd ventricle. **c**, Population Ca^2+^ activity in SCN^NMS^ neurons during several sleep-wake cycles. **d**, Ca^2+^ activity averaged over wake (55 episodes per mouse), NREM sleep (55 episodes per mouse) and REM sleep (12 episodes from each mouse) episodes from 3 mice. Mean ± SEM, one-way ANOVA (F_(2,363)_ = 79.26, *p* < 0.0001), followed by Tukey’s multiple comparison test. **** *p* < 0.0001. **e**, Ca^2+^ activity in SCN^NMS^ neurons at arousal state transitions. Upper panels: mean (± SEM) Ca^2+^ activity across all transitions (n = 3 mice). Lower: heatmaps depicting individual arousal state transitions from NREM sleep to wake (11 transitions per mouse), wake to NREM sleep (22 transitions per mouse), NREM sleep to REM sleep (11 transitions per mouse) and REM sleep to wake (9 transitions per mouse). Black dotted line indicates time of transition. Paired two-tailed t-test. ***p < 0.001; **** *p* < 0.0001. **f**, Mean (± SEM) Ca^2+^ activity over all mice at the initiation of the light pulse. Black dotted line indicates onset of the light pulse. Paired two-tailed t-test. **p < 0.01. Heatmap depicting Ca^2+^ activity at the light pulse for each mouse. White dotted line indicates onset of the light pulse.

**Extended Data Fig. 4.**
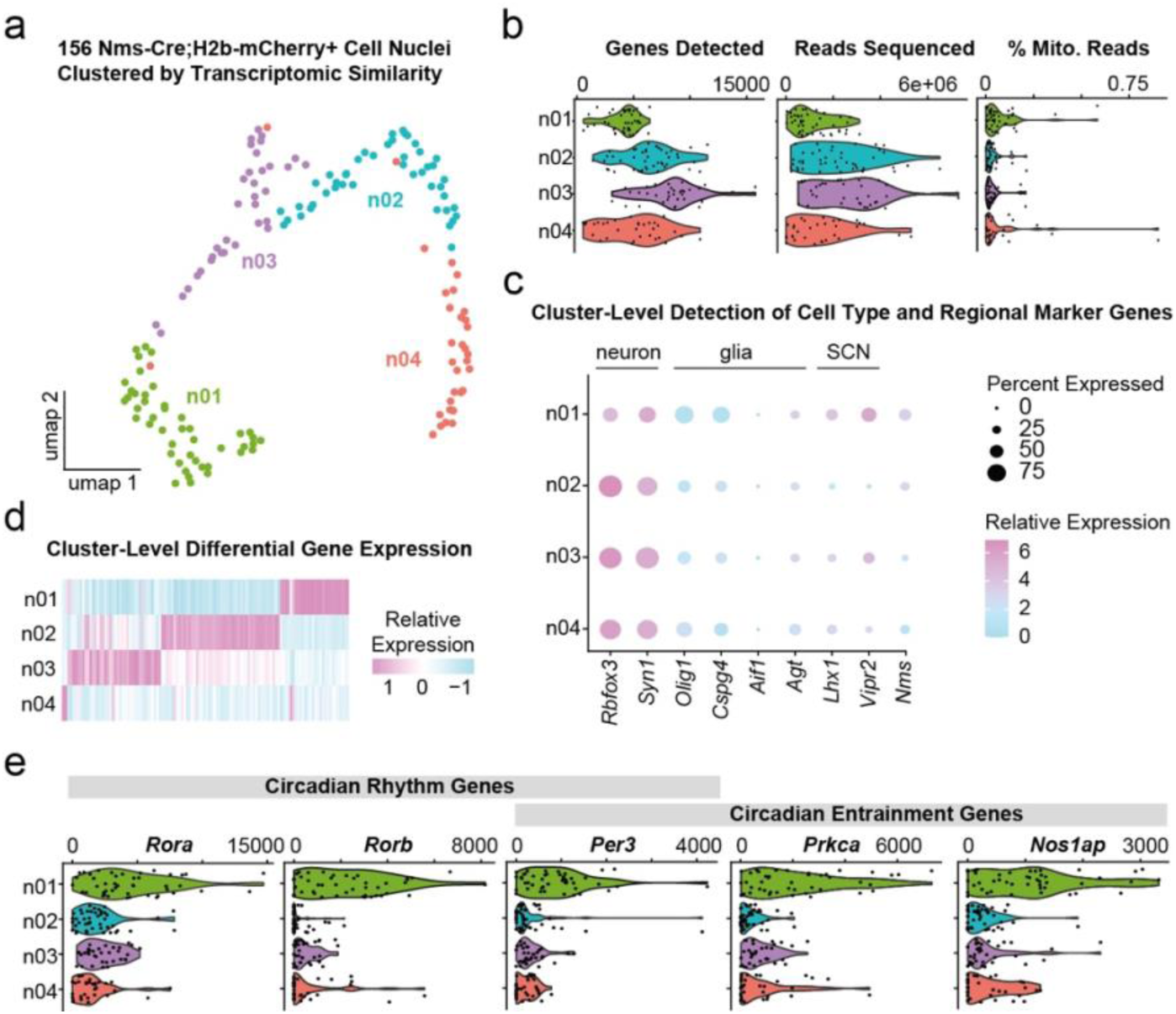
Molecular subtyping of *Nms*+ neurons in the SCN. **a**, Uniform Manifold Approximation and Projection (UMAP) two-dimensional embedding of single-nuclei transcriptomes after Louvain clustering in high-dimensional gene space. **b**, Violin plots of genes detected per nucleus, reads sequenced per nucleus, and percentage of reads from mitochondrial genes. **c**, Dot plot of cluster-level detection rate and expression of select marker genes. **d**, Heatmap of genes differentially expressed across clusters. **e**, Violin plots of circadian rhythm and entrainment genes from KEGG analysis of differentially expressed genes in cluster n01.

